# In-cell Structure and Variability of Pyrenoid Rubisco

**DOI:** 10.1101/2025.02.27.640608

**Authors:** Nadav Elad, Zhen Hou, Maud Dumoux, Alireza Ramezani, Juan R. Perilla, Peijun Zhang

**Author notes:** Equal contribution (Nadav Elad, Zhen Hou).

## Abstract

Ribulose-1,5-bisphosphate carboxylase/oxygenase (Rubisco) is a key enzyme in the global carbon cycle, catalyzing CO_2_ fixation during photosynthesis. To overcome Rubisco’s inherent catalytic inefficiency, many photosynthetic organisms have evolved CO_2_-concentrating mechanisms. Central to these mechanisms is the pyrenoid, a protein-dense organelle within the chloroplast of eukaryotic algae, which increases the local concentration of CO_2_ around Rubisco and thereby enhances its catalytic efficiency. Although the structure of Rubisco has been extensively studied by *in vitro* methods such as X-ray crystallography and single particle cryo-EM, its native structure within the pyrenoid, its dynamics, and its association with binding partners remain elusive. Here, we investigate the structure of native pyrenoid Rubisco inside the green alga *Chlamydomonas reinhardtii* by applying cryo-electron tomography (cryo-ET) on cryo-focused ion beam (cryo-FIB) milled cells, followed by subtomogram averaging and 3D classification. Reconstruction at sub-nanometer resolution allowed accurate modeling and determination of a closed (activated) Rubisco conformation. Comparison to other reconstructed subsets revealed local variations at the complex active site and at the large subunit dimers interface, as well as association with binding proteins. The different structural subsets distribute stochastically within the pyrenoid. Taken together, these findings offer a comprehensive description of the structure, dynamics, and functional organization of Rubisco within the pyrenoid, providing valuable insights into its critical role in CO_2_ fixation.

## Introduction

In the effort to mitigate global warming, carbon fixation, as a part of photosynthesis, has come into the spotlight^1^. Additionally, the ever-growing global population places a formidable burden on the sustainable food supply, which is closely linked to the efficiency of carbon fixation^1–4^. One-third of the global CO_2_ is arguably fixed in algae by ribulose-1,5-bisphosphate carboxylase-oxygenase, commonly known as Rubisco^5^. As the most abundant protein, Rubisco is indispensable for life on Earth as it catalyzes the reaction between CO_2_ and ribulose-1,5-bisphosphate (RuBP), resulting in the production of two molecules of 3-phosphoglycerate (3-PGA) in the Calvin-Benson cycle^6–8^. Despite its crucial role, Rubisco is known for its relatively slow catalytic rate and its dual activity, meaning it can also react with O_2_ instead of CO_2_, which in turn reduces the efficiency of carbon fixation^2,8–11^.

The structure of Rubisco is highly conserved across different species, reflecting its essential role in carbon fixation^2^. Rubisco is a holoenzyme, typically composed of two types of subunits. In higher plants, algae, and cyanobacteria, the enzyme is a hexadecamer, consisting of eight large subunits and eight small subunits^7,8,12–16^. The large subunits form the core of the enzyme where the catalytic sites are located, while the small subunits play a regulatory role, influencing the enzyme’s activity and stability^7,17^. Numerous structures of purified Rubisco have revealed the catalytic cycle in detail. The active site is located at the interface between the C-terminal of one large subunit and the N-terminal of an adjacent one, such that two large subunits form a functional dimer harboring two active sites. A rigid framework is formed around the active site, while several mobile loops, including the N and C termini and the important loop 6, mediate the opening and closing of the substrates binding pocket^7,18–20^. Yet, it remains to be seen whether the structural patterns and dynamics that have been revealed *in vitro*, are maintained in the crowded environment of the cell, and how the cellular Rubisco population varies with respect to the catalytic activity.

In the unicellular green alga *Chlamydomonas reinhardtii*, Rubisco is sequestered within a specialized non-membrane-bound organelle called the pyrenoid, located in the chloroplast^21–23^. The pyrenoid is primarily found in many eukaryotic algae and plays a crucial role in enhancing the efficiency of carbon fixation, a process believed to have been driven by the gradual decrease of atmospheric CO_2_ over billions of years until the recent industrial revolution by humans^21–25^. This compartmentalization is particularly advantageous under low CO_2_ conditions, as it helps to concentrate CO_2_ around Rubisco, minimizing the enzyme’s oxygenase activity and reducing photorespiration^26–29^.

The concentration of Rubisco in the pyrenoid of *C. reinhardtii* is mediated by the protein Essential Pyrenoid Component 1 (EPYC1), forming liquid-liquid phase separation (LLPS)^13,30,31^. Unlike its analogues in cyanobacteria: the linker protein CsoS2 in α-carboxysomes and CcmM in β-carboxysomes which bind to the interface of two large subunits of Rubisco^14,15,32^, EPYC1 binds to the small subunit directly^13^. The difference in binding sites on Rubisco by these linker proteins is believed to be related to the packaging of Rubisco. As the packaging of Rubisco affects the utilization and assimilation of CO_2_, understanding the *in-situ* organization of Rubisco particles has garnered significant attention. Recent studies have demonstrated intriguing packaging patterns for Rubisco in α- and β-carboxysomes which are confined by shells^15,33,34^. Whether similar or distinguished packaging patterns exist for Rubisco in the pyrenoid remains elusive. Pioneering in-cell cryo-electron tomography (cryo-ET) works on the pyrenoid in *C. reinhardtii* have revealed its tight connection with thylakoids and presented a single in-cell structure of Rubisco at 16.5 Å, suggesting potential packing patterns for Rubisco^13,21,35^. However, higher-resolution in-cell structures of Rubisco and more detailed spatial analyses of Rubisco in the pyrenoid are essential to build a robust and comprehensive model that fully captures the complexity and dynamic nature of this organelle and its critical role in life-sustaining processes.

In this work, we employed cryo-focused ion beam (cryo-FIB) milling and cryo-ET, coupled with subtomogram averaging (STA) to investigate the pyrenoid and Rubisco in *C. reinhardtii*. We obtained an in-cell structure of Rubisco at sub-nanometer resolution, enabling us to perform the molecular dynamic flexible fitting (MDFF), thereby revealing a closed conformation of Rubisco in the cell. Successive in-cell analyses show the localized heterogeneity of Rubisco conformations and pyrenoid binding partners, reflecting its dynamic nature and enzymatic activity in carbon fixation. Moreover, the distribution of Rubisco in pyrenoid is revealed to be stochastic, different from that rigid packaging in the α- and β-carboxysomes. Taken together, our work enhances the understanding of the pyrenoid and Rubisco in intact cells, providing valuable insights for future studies on fundamental mechanisms and the bioengineering of pyrenoid and Rubisco in other organisms.

## Results

### Structural heterogeneity of Pyrenoid Rubisco is revealed by the classification of sub-volumes

To gain insight into the structure of pyrenoid Rubisco, we vitrified *C. reinhardtii* cells on TEM grids and used cryo-FIB to cut thin lamellae through the cells. We then applied cryo-ET to collect tilt series from lamellae at pyrenoid areas (Supplementary Tables 1,2). Rubisco complexes are highly abundant within the pyrenoid and appear in their expected spherical morphology (Fig. 1a). Additionally, thylakoid and pyrenoid tubules can be clearly seen (Fig. 1a-c). To resolve the structure of Rubisco, deep learning-based particle picking was performed, and most discernable Rubisco particles were correctly picked (Fig. 1c). In the following STA analysis, iterative 3D classification and refinements were applied. Interestingly, even though Rubisco particles were ubiquitously present in tomograms and seemed to have similar overall morphology, iterative 3D classifications revealed vast heterogeneity at medium-low resolutions (Supplementary Fig. 1, Supplementary Table 3, and see below). Experimenting with different 3D classification strategies, we found that classification and cleaning the data from “junk” particles without applying symmetry gave more reliable results compared to applying D4 symmetry from the start. “Good” 3D classes were selected based on the overall appearance of Rubisco features, the map’s resolution, and accuracy in the angular assignment. Following classification with no applied symmetry, classes were refined separately with D4 symmetry applied and finer bin sizes to resolutions of 8 to 15 Å. We start by describing the best-resolved Rubisco class and the associated secondary structure model.

**Fig. 1:**
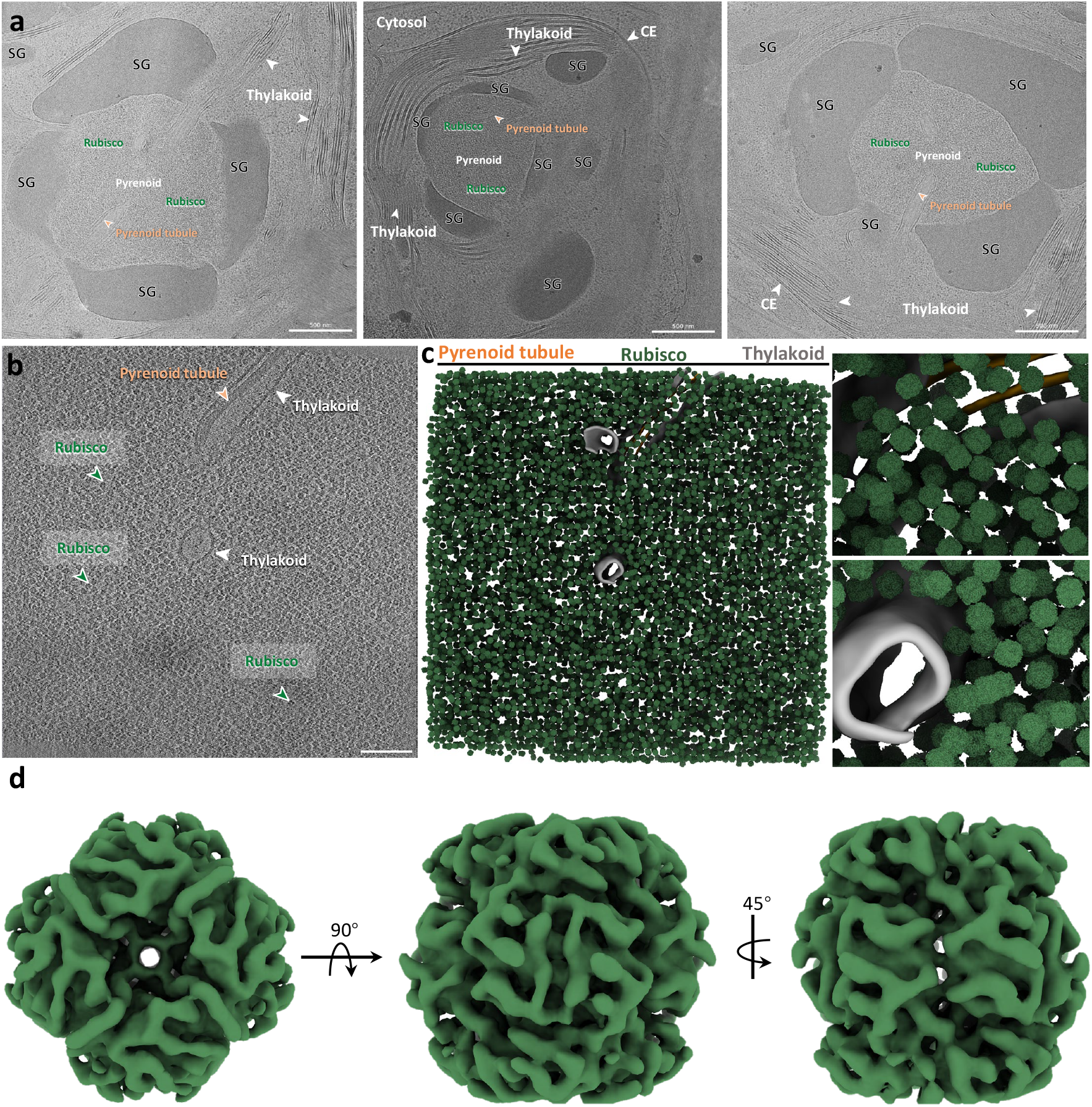
Cryo-ET of pyrenoid and in-cell structure of Rubisco. **a** overviews of phase-separated Rubisco in intact pyrenoids of various sizes. Cellular components are labelled and annotated accordingly, SG: starch granule, CE: chloroplast envelope. **b** A representative tomographic slice of the pyrenoid in *C. reinhardtii*. Thylakoid, pyrenoid tubule, and Rubisco particles are labeled and indicated accordingly. Scale bar = 100 nm. **c** The segmented volume of the tomogram in **b**. The left shows the overview of the segmented pyrenoid with mapped-back Rubisco particles, the right showcases two close-up views of Rubisco particles with the thylakoid. Thylakoid, pyrenoid tubule, and Rubisco are coloured accordingly. Rubisco model is adapted from the in-cell structure of this work. **d** The in-cell structure of Rubisco in *C. reinhardtii* shown in three orthogonal views (from *n* = 17,713, a subclass from *n* = 198,692, Number of tomograms = 26).

### The best-resolved Rubisco complex is in a closed conformation state

The best resolved 3D class (number 12) was refined to 8.1 Å resolution, enabling the identification of secondary structure elements (Fig. 1d, Supplementary Fig. 2). To our knowledge, this is the highest resolution structure of Rubisco, reconstructed directly from the cellular environment without purification. The map contains 12% of the good particles (i.e., 148,639 particles contained in the C1-refined classes, Supplementary Fig. 1b, Supplementary Table 3). Docking in the published X-ray coordinates from purified *C. reinhardtii* Rubisco complexes revealed extensive similarity between our *in-situ* structure and the published purified ones (Fig. 2a). All the α-helices and β-sheets can be accounted for in the map, while most loops can be located, indicating the map’s quality.

**Fig. 2:**
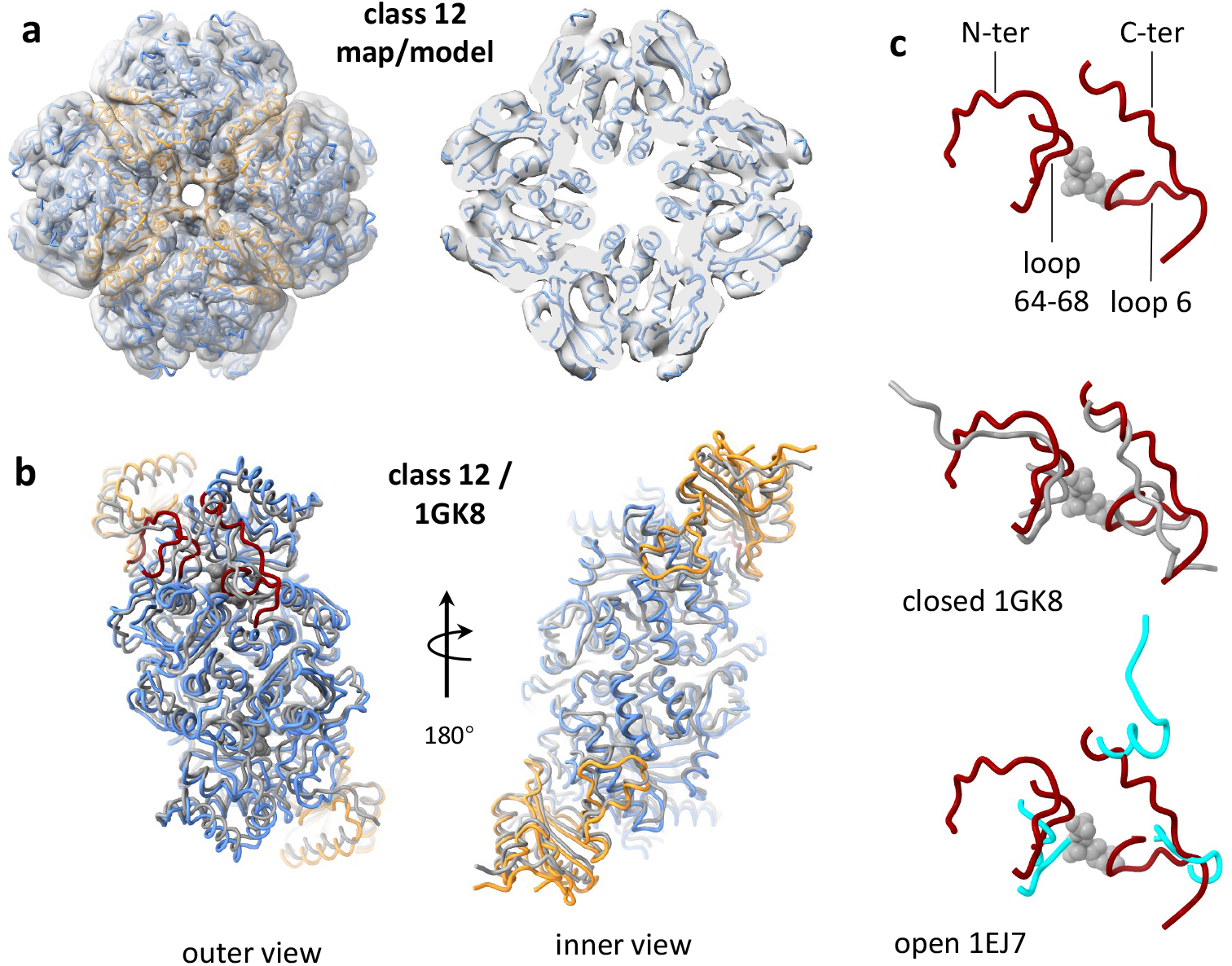
Structure of best-resolved STA class. **a** Map and fitted MDFF model of class 12, which refined to the highest resolution (8.1 Å). Shown are top and cut-away views. Large and small subunits are colored blue and orange, respectively. **b** Alignment between two large subunits and two small subunits of class 12 MDFF model (light blue and orange, respectively) and PDB 1GK8 (grey). Key active site segments in the MDFF model are colored dark red. Shown are LS Loop 6 (aa 331-338), C-terminus (aa 461-475), loop 64-68, and the resolved N-terminus (aa 7-21). The CABP inhibitor from 1GK8, which marks the position of the active site, is shown in atomic view. **c** Comparison of active site conformations. The conformational changes in these residues correlates with the Rubisco activity state. The MDFF model (dark red) is compared to PDB 1GK8 (grey, middle panel) and PDB 1EJ7 (cyan, bottom panel), which represent closed and open conformations, respectively. A better match is found for the closed state. The CABP inhibitor from 1GK8 is shown in atomic view in all panels for reference.

To better understand the reconstructed conformational state, we used molecular dynamics flexible fitting (MDFF, detailed in the methods section) to refine the published Rubisco coordinates within our map (Fig. 2a). Even though side chains cannot be resolved at this resolution, secondary structure elements from the x-ray structures can be modeled reliably, in conjunction with known protein structural constraints. We then compared the refined coordinates to published structures of known functional states. Overlaying the class 12 MDFF coordinates with those of PDB 1GK8^16^ resulted in a root-mean-square distance (RMSD) of 2.75 Å. An excellent match is seen at the center of the complex near the 2-fold axis and at the interior-facing parts of the large subunit (Fig. 2b). The MDFF structure is more extended along the 4-fold axis compared to the 1GK8 structure, producing mismatches of up to about 4 Å between corresponding helices at the small subunits and nearby large subunit segments.

However, rather than major whole-domain movements, the transition between open (inactivated) and closed (activated) states has been shown to involve significant local conformational changes of loops around the active site. In particular, the large subunit loop 6 (aa 331-338) and C-terminal (aa 461-475) cover the active site in the closed conformation, and both fold outwards and become disordered in the open conformation. Loop 64-68 and the large subunit N-terminal (aa 7-21) cover the other side of the active site in the closed conformation and are disordered in the open conformation. We compared these elements between class 12 MDFF model and either 1GK8^16^ or 1EJ7^20^ coordinates, representing a fully closed or fully open conformation, respectively (Fig. 2c). The conformations of these four elements in our model clearly match the closed-state conformation as they overlay well with the 1GK8 structure. Most strikingly, the positions of mobile loops 6 and 46-68 clearly cover the location of the active site, and map density is observed at this position up to high contour values (Supplementary Fig. 3a-c). The map features density which correlates well with the expected closed conformations of the C and N termini, although these are fully present at lower sigma values. This may indicate a higher level of heterogeneity or can be due to their peripheral locations. As seen in Supplementary Fig. 3d-f, the N terminus is well accounted for in the map, while the C-terminus density is interrupted in the middle, around Glu468. Indeed, the C-terminus is known to be poorly anchored to the body of the large subunit even in the closed conformation, except for a network of interactions formed at the “latch site” by the conserved Asp437. This interaction stabilizes the closed active site and is important for catalytic efficiency and specificity^20,36^. The MDFF model positioned the end of the C-terminus in contact with the body of the complex within a strong peripheral density, correlating well with the location of the latch site (Supplementary Fig. 3e, f). This density is therefore more likely to be part of the C-terminus than a binding protein.

### Local conformational changes and binding proteins are the main sources of heterogeneity

Processing of a large dataset of pyrenoid Rubisco subtomograms has revealed the presence of multiple distinct classes of complexes (Supplementary Fig. 1). Exploring the origins of this heterogeneity is particularly intriguing, as the dataset captures authentic frozen snapshots of functional Rubisco, representing its dynamics and interactions with binding partners in the cell. Varying sources of heterogeneity can exist in such a cellular environment, including concerted whole-domain movements, local conformational variations, disruption of symmetry, and the presence of binding proteins. Yet, significant heterogeneity essentially limits the attainable resolution because it compromises the number of particles per class and, consequently, reduces the signal-to-noise ratio (SNR) in the average map.

We first compared the extent of whole domain movements between classes. To this end, we fitted the Rubisco coordinates into the three D4-symmetric classes that were refined to the highest resolution (classes 7, 10, 13). The Class 12 MDFF model (described above, Fig. 2a) was used as a starting model for rigid body fitting of whole domains, and as a reference for comparison. As seen in Supplementary Fig. 4, only subtle whole-domain shifts of up to 2.5 Å occur between the class 12 model and the models fitted into all three classes. Although the extent of the observed movements is much smaller than the resolution of the maps, this result is in accordance with x-ray structures showing that Rubisco activity cycle and varying conditions induce only minor, angstrom-scale, whole-domain movement^7,19,20^.

Next, we explored local variations in the maps as a source of heterogeneity between classes. Some local variations can be readily observed when comparing maps by visual inspection (Fig. 3a), however, they can be better tracked and highlighted by subtracting maps from one another and looking at the resulting difference maps (Fig. 3b-g). To this end, we calculated difference maps from confidence maps binarized at 1% false discovery rate threshold^37^. Interestingly, significant differences (density gain) occur between class 12 and all other classes at the active site (Fig. 3b-d blue). As indicated above (Fig. 2c) and shown before^18,38^, the four large subunit segments - loop 6, the N- and C-termini, and loop 64-68, are particularly mobile during the catalytic cycle, shifting between a closed/active state and an open/inactive state, as well as populating intermediate states. The observation of significant density variation at this location suggests that the pyrenoid Rubisco population exists in multiple activity states. Class 12, which is in the closed state, is particularly different from the rest of the classes at this position and perhaps constitutes a more stable conformation (Fig. 3b-d). Smaller-scale differences at the active site can be seen between the other three classes (Fig. 3e-g).

**Fig. 3:**
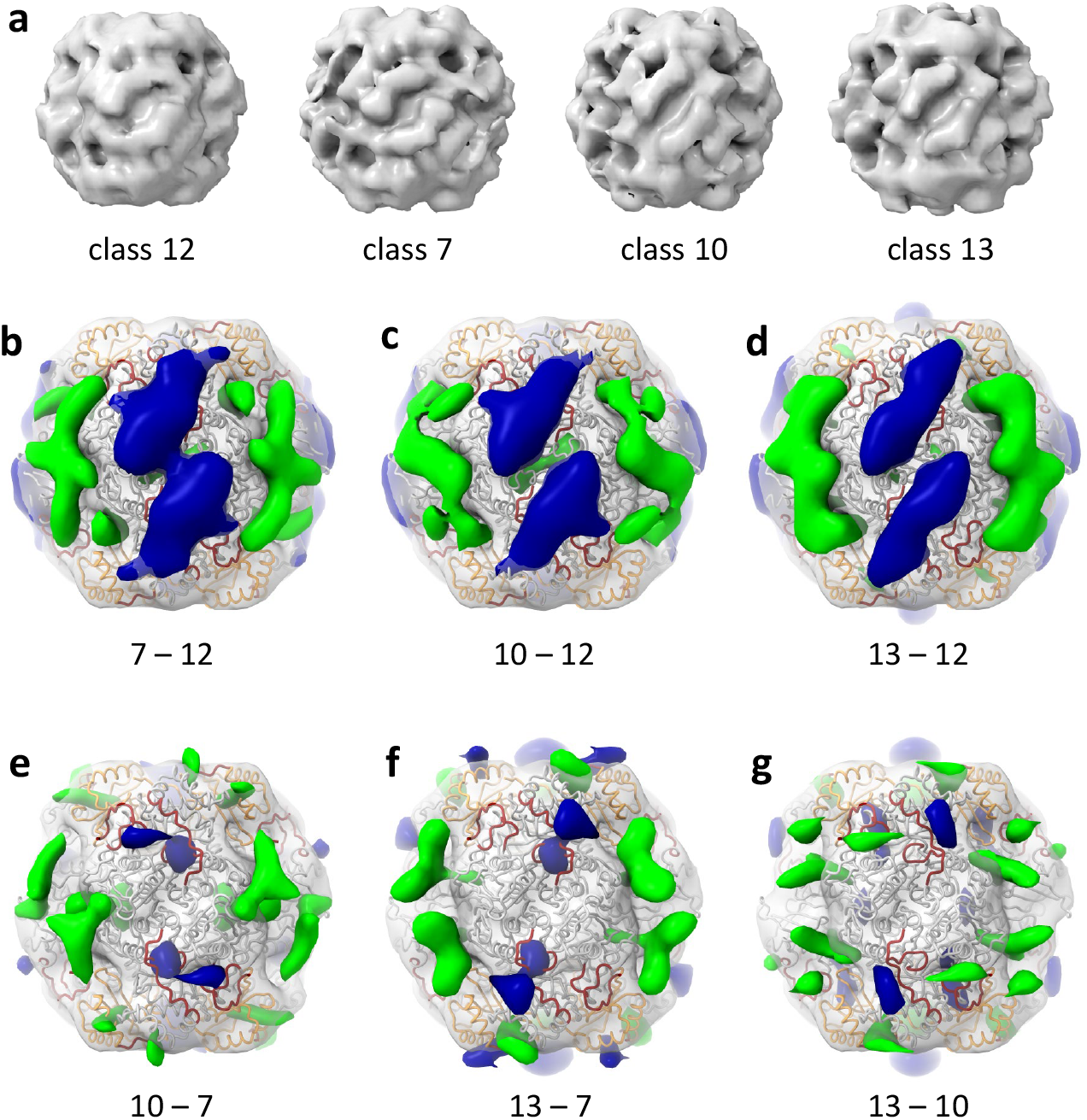
Structural heterogeneity of pyrenoid Rubisco. **a-c** D4-symmetric maps of four classes out of ten (Supplementary Fig. 1), which were refined to the highest resolution. Class 12 (8.1 Å) and 7 (12.3 Å) maps were low-pass filtered to 13.6 Å to match the resolution of classes 10 and 13. **b-g** Difference maps showing major variations in density between classes. Positive densities resulting from the subtraction of one map from another as indicated in the panels are colored dark blue and negative densities are colored green. Difference maps between classes were calculated from confidence map binarized at 1% false discovery rate threshold, gaussian filtered, and presented at 10 SDs contour level. For orientation, the difference maps are overlayed on the class 12 low-passed filtered map and fitted MDFF model (Fig. 2a). Large subunits, small subunits and active site residues are colored grey, orange, and dark red, respectively.

Significant conformational variability is seen also between the large subunit dimers near the 2-fold axis. As observed for the active site, class 12 shows major differences (density loss) at this interface compared to the other 3 classes (Fig. 3b-d, green). This difference density can be interpreted as correlated movement of subunits at the dimer interface. Additionally, significant difference densities protruding outside the Rubisco model appear next to the large subunit dimer interface when comparing classes 7, 10 and 13 (Fig. 3e-g, green). This position correlates well with the binding interface for carboxysome matrix proteins^14,15,32,34,39,40^, and likely indicates the presence of Rubosco binding proteins within the pyrenoid. Finally, difference densities are observed at several locations adjacent to the small subunits (Fig. 3e-g), including at the opening of the cavity on the 4-fold axis (Fig. 3d,f,g blue).

### Asymmetric binding to Rubisco

Functional pyrenoid Rubisco comprises 8 identical copies of the large and small subunits arranged in D4 symmetry. However, the Rubisco binding proteins may not bind to all subunits simultaneously^31,35^, and conformational symmetry is not essentially maintained in the cellular context^41^. To explore the notion of asymmetricity in Pyrenoid Rubisco, we refined the sub-volume classes without applying D4 symmetry (Supplementary Fig. 1b, Supplementary Table 3). These C1 refinements compromised resolution compared to the corresponding D4 symmetric reconstructions. In particular, the best-resolved class 12 map was now refined to 13.1 Å, making the analysis of inter-subunit conformational heterogeneity impractical. Interestingly however, we observed that the C1 class 12 map presents a number of extra densities at the expected location of the EPYC1 binding site, next to the small subunits (Fig. 4a). EPYC1 was previously shown to have multiple alpha-helical repeats, which bind to the Rubisco small subunit, flanked by unstructured spacer regions^13,31^. Simultaneous binding of multiple Rubisco complexes supports its clustering in the pyrenoid.

**Fig. 4:**
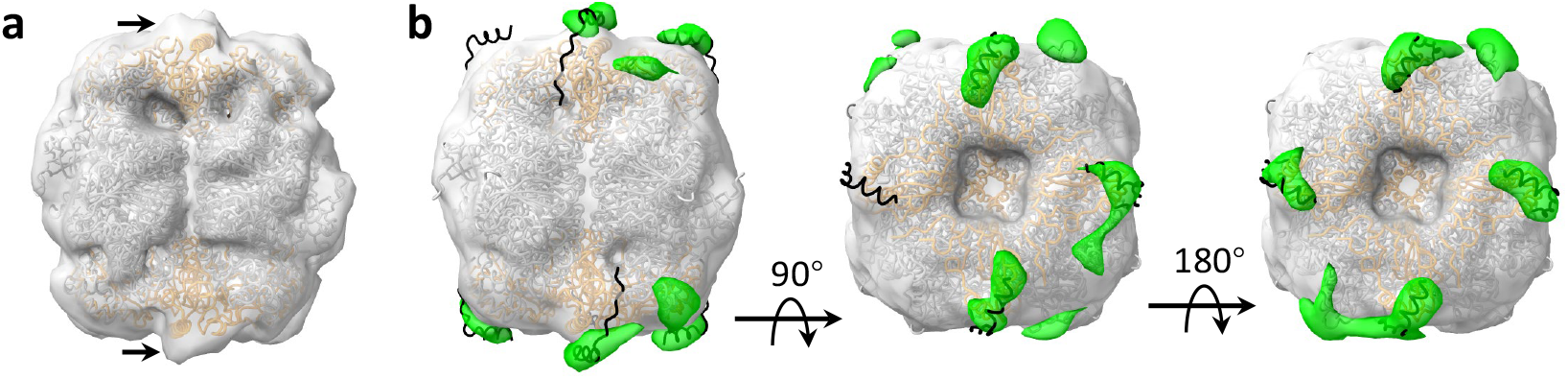
Asymmetric binding of EPYC1 to Rubisco. **a** Asymmetric map of class 12 with. The MDFF model from the D4 symmetric class 12 map was fitted as a rigid body (large and small subunits are colored grey and orange, respectively). Arrows indicate extra density next to the small subunits. **b** Difference density between the C1 and D4 symmetric class 12 maps (green), overlayed on the D4 symmetric class 12 map (low-passed filtered, transparent white). The difference map was calculated from confidence map binarized at 1% false discovery rate threshold, gaussian filtered, and presented at 13 SDs contour level. The SPA structure of Rubisco-EPYC1 (PDB 7JFO) was fitted as a rigid body into the map. The difference densities varies in shape and matches the location of most EPYC1 helices (black coordinates). The difference density was calculated from confidence maps, binarized at 1% false discovery rate threshold.

The potential EPYC1-assiciated extra densities were not observed in the class 12 D4 symmetric map, likely because they were averaged out when applying symmetry. Thus, to validate the extra densities seen in the class 12 C1 reconstruction, we calculated a difference map between C1 and D4 maps, binarized at 1% false discovery rate threshold. As can be seen in Fig. 4b, the most significant difference in densities can be associated with the EPYC1 helix, since they are located next to the small subunits at the predicted binding site. These densities vary in shape and are absent next to one of the small subunits, indicating partial occupancy.

### Stochastic distribution of Rubisco in the pyrenoid

To investigate whether Rubisco particles are organized in a specific pattern within the pyrenoid as seen for Rubisco in α- and β-carboxysomes, we first analyzed the distances and angles between the nearest neighbouring particles in class 12. The results revealed a largely random distribution, with a mean paired distance of ∼13 nm (Fig. 5a) as previously observed^35^. This is consistent with the measured paired distances of 12.8 nm and 12.9 nm in two α-carboxysomes^15^, but slightly larger than the distance in β-carboxysomes (122 nm)^34^. The pairwise average angle was measured ∼90° (Fig. 5b). To explore potential patterns among Rubisco particles with restricted paired distances and angles, we categorized particles based on these parameters and mapped each category back to the tomograms. While no distinct global patterns emerged (Fig. 5b, c, Supplementary Fig. 5-7), some local clusters were identifiable under stringent distance and angle constraints (Fig. 5d, e).

**Fig. 5:**
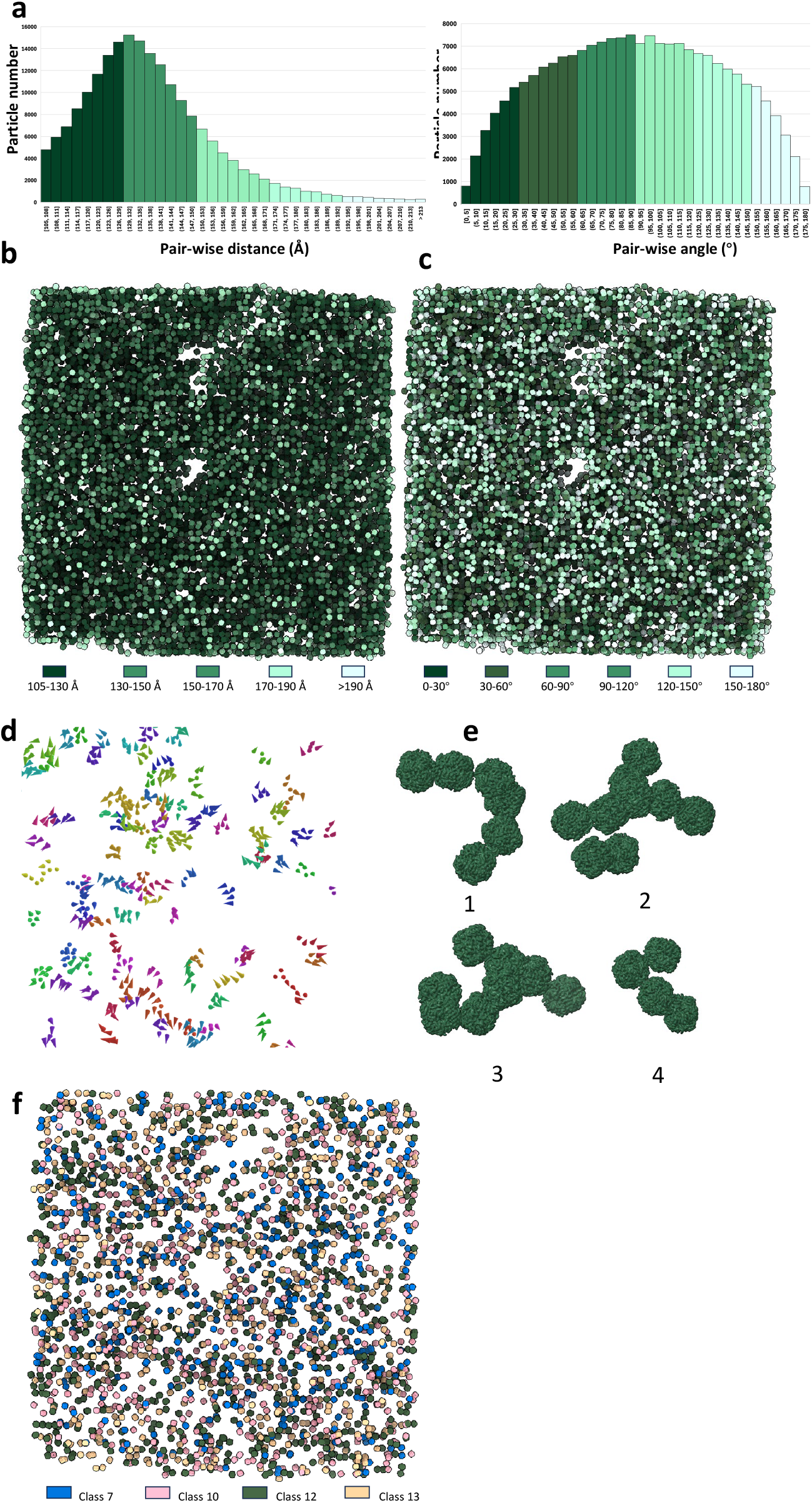
Mapping-back of Rubisco in the pyrenoid. **a** Distributions of pair-wise distances and angles between the nearest Rubisco neighbours (from *n* = 215,272, *n* of tomograms = 26). **b** Mapping-back of Rubisco by pair-wise distance of nearest neighbours, Rubisco is coloured in various shades of green accordingly. **c** Mapping-back of Rubisco by pair-wise angle of nearest neighbours. Rubisco is coloured in various shades of green accordingly. **d** Local clustering of Rubisco with pair-wise distances ranging from 135 to 165 Å, pair-wise angles from 0° to 40, and minimum cluster forming particle number = 5. Various local clusters are coloured accordingly, and the pointed end of cones indicate the orientation of particles. **e** close-up look of four representative local clusters of Rubisco. **f** Mapping-back of Rubisco by classes, with four major classes included and coloured accordingly. The mapping back is exemplified with the tomogram in **Fig. 1b**.

Given that STA analysis identified multiple classes of Rubisco in the pyrenoid, reflecting different states, we examined whether specific classes displayed preferential localization. Mapping the four best-resolved classes back to the tomograms showed no evidence of class-specific clustering (Fig. 5f). In summary, Rubisco particles in the pyrenoid exhibit a predominantly stochastic distribution, with evidence of local clustering of some particles in the pyrenoid.

## Discussion

In the present study, we provide a detailed structural analysis of the Rubisco complex as it exists within the native pyrenoid. While the structure of Rubisco and its catalytic cycle have been thoroughly investigated *in vitro* using x-ray crystallography and single particle cryo-EM, these studies primarily relied on purified samples and focused on a single stable conformational state. Here, we employed cryo-FIB milling of *C. reinhardtii* pyrenoid sites in combination with cryo-ET, subtomogram averaging, and detailed 3D classification, allowing us to capture snapshots of functional Rubisco complexes in their native cellular environment, thereby avoiding the artifacts that can be involved in purification.

The classification of a large subtomogram dataset revealed multiple conformations of the complex, along with external densities, which can be interpreted as partial structures of binding proteins. The identification of these diverse structural patterns indicates that Rubisco complexes exhibit variations in their activity states and binding partners within the pyrenoid matrix. The different classes do not show any specific spatial localization within the pyrenoid, nor do they originate from a particular cell (lamellae), rather, they are randomly distributed. Consequently, we find no evidence for synchronization or compartmentalization of Rubisco dynamics.

The vast heterogeneity present in the sample limits the achievable resolution, as it imposes classification into small subsets. Nevertheless, one of the classes was resolved at a resolution of 8.1 Å, allowing for accurate modeling of secondary structures. MDFF modeling indicated that this class features a Rubisco population in a closed conformation state, while comparison to other, lower resolution classes exposed the undelaying structural heterogeneity. Our study confirms that Rubisco does not exhibit significant whole-domain movements within the pyrenoid, consistent with findings from purified complexes^7^. The main variations between classes are characterized by local conformational changes, primarily around the active sites and the interface between large subunit dimers, while other significant variations are associated with locations and morphologies of binding proteins. The cores of the large and small subunits remain rigid, devoid of significant local conformational changes. Given this core rigidity, it is intriguing to consider the factors that limit resolution in this preparation. We suggest that several factors may contribute, including the high density of complexes within the pyrenoid, the gel-like matrix of unstructured proteins surrounding the Rubisco complexes, and sample thickness, all of which reduce SNR in the tomograms. Other factors that limit resolution may include asymmetries in catalysis between subunits^41^ and variations in binding protein interactions, which were partially elucidated here.

We observe evidence of Rubisco binding proteins primarily at two interfaces: adjacent to the small subunits and the at interface between large subunit dimers. The former has been shown to be the interface of the EPYC1 linker protein or other pyrenoid proteins which share a similar binding motif^13,42^. The latter is the binding interface of CsoS2 or CcmM linker proteins in α- or β-carboxysome Rubisco, respectively^14,15,32,34,39^, as well as for the carbonic anhydrase enzyme, CsoSCA^40^. It is yet unclear which proteins bind to the pyrenoid Rubisco at the large subunit dimer interface and what their functions may be. Interestingly, the density for EPYC1 is particularly evident in the best-resolved asymmetric map, and is averaged out in the D4 symmetric maps, suggesting partial occupancy on the small Rubisco subunits. In contrast, the density associated with the large subunit dimer interface is evident in the D4 symmetrized maps and appears at a lower resolution. This indicates that a subset of the Rubisco population exhibits a high occupancy of these binding proteins.

In summary, this study offers direct observations of the structure and function of Rubisco within the native and intact pyrenoid, enhancing our understanding of the carbon fixation mechanism in algae. Additionally, the study proposes a framework for further investigations of native Rubisco at higher resolutions.

## Methods

### Strains and cell culture

*C. reinhardtii* wild-type strain CC-1690 was kindly given by Prof. Dr. Luning Liu at the University of Liverpool. Cells were maintained in Tris-acetate-phosphate (TAP) medium (Thermo-Fisher Scientific) under a normal circadian cycle (12-hour light and 12-hour dark) with the light intensity at 12,000 lx at room temperature on a shaker at 150 rpm.

### Plunge-freezing vitrification

An aliquot of 3.5 μl *C. reinhardtii* cell suspension at 1.5×10^6^ cells/ml was applied to the glow-discharged holey carbon-coated copper grid (R 2/1, 200 mesh) (Quantifoil) and blotted on the back of the grid for 9 seconds by Leica GP2 (Leica Microsystems), followed by the plunge freezing in liquid ethane.

### Cryo-FIB milling

Lamellae were prepared using an Aquilos 2 cryo-FIB/SEM (Thermo Fisher Scientific) located at electron Bio-Imaging Centre (eBIC), UK or Arctis plasma cryo-FIB/SEM (Thermo Fisher Scientific) located at the Rosalind Franklin Institute, UK (Supplementary Table 1).

Lamellae preparation on Aquilos 2: Thinning was performed on a rotatable cryo-stage maintained at −191 °C via an open nitrogen circuit. Prior to milling using a Gallium beam at 30 kV, grids were mounted onto a shuttle, transferred to the cryo-stage, and coated with an trimethyl(methylcyclopentadienyl) platinum (IV) (organometallic platinum) layer using the GIS system (Thermo Fisher Scientific) for 30 seconds. Cells located near the centers of grid squares were selected for thinning. The process was carried out in a stepwise manner using the automated milling software AutoTEM software (Thermo Fisher Scientific), with currents decreasing incrementally from 0.5 nA to 30 pA at 30 kV. The final thickness of the lamellae was set to 120 nm.

Lamellae preparation on Arctis: Autogrid clipped TEM grids were loaded chamber via the robotic delivery device (Autoloader) in a dual-beam plasma focused ion beam scanning electron microscope with redesigned sample, which is a prototype for the commercially available Arctis microscope (Thermo-Fisher Scientific). The ion species used for milling was Argon at 30kV. Both ion and SEM columns are aligned, and the measured currents are within 20% of their target. SEM imaging was done on the grids to ensure suitability for lamella preparation. Once the sample is selected, a conductive layer has been deposited on the sample by milling a platinum target on the sample (16kV 1.4 μA). Then a protective layer of organo-platinumwas deposited using a gas injection system. Finally, another conductive layer of platinum was deposited. Lamella sites were identified and queued in the AutoTEM software (ThermoFisher Scientific) beforeunattended milling using the following sequence: i) stress relief trenches (0.74 nA) ii) three steps coarse milling (2, 0.74 and 0.2 nA) iii) two steps polishing (60 pA and 20 pA).

### Cryo-electron tomography data collection

The tilt series were acquired using two Titan Krios G4 TEMs (Thermo-Fisher Scientific) with similar setups, one at eBIC and one at RFI (Supplementary Table 2). Both microscopes were operated at 300 kV with fringe-free illumination. Imaging was done on a Falcon 4i direct detector installed behind a Selectris X energy filter (Thermo-Fisher Scientific), using a slit of 10 eV. All tilt series were recorded at a nominal magnification of 64,000×, corresponding to physical pixel sizes of 1.97 Å (eBIC) or 1.978 (RFI), using the dose-symmetric scheme starting from the lamella pre-tilt of −12° or 12° (dependent on grid orientation), 2° increments and tilt span of 54°. Tilt series were acquired using an automated low-dose procedure implemented in Tomography 5 software (Thermo-Fisher Scientific). In some positions multiple tilt series were acquired using image-beam shift. The nominal defocus range was 2.5 to 4.5 µm, and the total dose was ∼120 e^-^/Å^2^. Each tilt series was fractionated into 6 movie frames.

### Subtomogram averaging

Movie frames were motion-corrected using MotionCor2^43^, followed by CTF estimation within individual tilts using CTFFIND4^44^ and tilt-series alignment and reconstruction in bin 2 (3.94 Å/pixel) using AreTomo^45^. In total 26 tilt-series were selected for analysis based on the quality of the reconstruction and appearance of extensive Rubisco clusters. Particle picking was done using crYOLO^46^. The network was trained by manually annotating 12 sections from 3 different tomograms with default parameters and 38-pixel boxes. Running prediction in tomography picking mode resulted in 515,948 subtomograms from all tomograms. Tilt-series, along with corresponding alignment files, CTF parameters, and particle coordinates, were imported into RELION4^47^ for running the STA pipeline. Subtomograms (RELION PseudoSubTomograms) were initially processed in bin 4 (7.88 Å/pixel) and 32^3^ box size. “Bad” subtomograms were discarded by two rounds of 3D classification into 10 classes, without applying symmetry. “Good” 3D classes were selected based on the overall appearance of Rubisco features, the map’s resolution, and accuracy in the angular assignment. The SPA structure of *C. reinhardtii* Rubisco filtered to 60 Å was used as an initial reference (PDB 7JN4^13^), and a mask of 180 Å diameter was applied during the process. The resulting 215,272 subtomograms were 3D classified into 20 classes with a mask of 160 Å diameter applied (Supplementary Fig. 1a). Refinement of all good classes was then performed at bin 2 (3.94 Å/pixel), box size 100^3^, with C1 or D4 symmetry applied, and mask diameters of 160 or 150 Å, respectively (Supplementary Fig. 1b, c, Supplementary Table 3). The best-resolved D4-symmetrized map (class 12, 17,713 subtomograms) was further refined at bin 1 (1.97 Å/pixel) and 256^3^ box size. CTF refinement (512^3^ box size for CTF estimation) followed by frame alignment and reconstruction with an applied solvent mask resulted in a final resolution of 8.1 Å (Supplementary Figs. 1d, 2).

Difference maps between classes (Fig. 3b-g) were prepared as follows: refined maps of classes 7 and 12 were low pass filtered to 13.6 Å to match the lower resolution of classes 10 and 13. Confidence maps were calculated with a noise box size of 27 pixels^37^, binarized at 1% false discovery rate threshold in ChimeraX^50^, and subtracted from each other. The resulting difference density was Gaussian filtered (SD=4) and displayed at a contour level of 10 SDs with “hide dust” set to 15 Å. The difference map in Fig. 4b was prepared in the same way, except that the low pass filtered, D4 symmetric class 12 map was subtracted from the corresponding C1 map.

### Real-space refinement of Rubisco coordinates within the cryo-EM map

A specialized real-space refinement protocol was developed for the present work (Supplementary Fig. 8). The protocol is detailed in the Supplementary Information and is summarized here. First, rigid-body docking was performed to embed the large subunits (PDB: 1GK8^16^) and small subunits (PDB: 1EJ7^20^) of the Rubisco structure into the best-resolved D4-symmetrized class 12 map. Mutations introduced during crystallization were reverted back to the wild-type sequence (UniProt ID P00877 for the large subunit and P00873 for the small subunit) using ChimeraX^48^. Flexible regions, namely loop 6, the N- and C-termini of the large subunit were generated de novo using Rosetta version 2024.09^49–52^. The C-terminus of the large subunit was restrained by maintaining the distances between D473-R134 and D473-H310, to guide the model into the cryo-EM density, as the interactions were observed in previous studies^20^. The model was further refined iteratively by applying a local rebuilding procedure using CartesianSampler mover in Rosetta. Protonation states of titratable groups were determined by using PD2PQR^53^, then ions and water molecules were added using VMD^54^. Molecular dynamics flexible fitting (MDFF) simulations^55,56^ were performed as described in Perilla et al^57^, while the backbone heavy atoms were coupled to the Cryo-EM density map using a grid-based biasing-potential^58^.

The MDFF-derived model revealed multiple solutions for the N- and C-termini coordinates of the large subunits, while in none of the chains both termini were fitted in the EM density simultaneously (Supplementary Fig. 9). Thus, a hybrid large subunit was created by combining residues 1-103, taken from the best-fitting N-terminus, and residues 104-475, taken from the best-fitting C-terminus chain. The complete Rubisco structure built from the hybrid large subunit model was subjected to further relaxation to remove possible steric clashes using NAMD3.0.1. The minimized-hybrid model was further refined iteratively by using an unsupervised^59^ and supervised protein refinement tool, applying FastRelax^60–62^ protocol in Rosetta on selected regions. The unsupervised refinement tool identified regions with low molProbity score^63,64^, while the supervised refinement tool selected regions having poor local-cross correlation (Supplementary Fig. 10). To improve the quality of the beta sheets, the beta strands coordinates on the large subunits were replaced by the coordinates of the corresponding residues in PDB: 1GK8 (Supplementary Fig. 11). Another round of symmetrical MDFF simulation were performed on unprotonated Rubisco structure. Lastly the model was subjected to minimization using NAMD3.0.1 without coupling the protein to the cryo-EM density. The resulting model was in good agreement with the density, as evidenced by the model to density FSC (Supplementary Fig. 12d), and the local cross-correlation (Supplementary Fig. 12e-f).

### Segmentation

To enhance the segmentation, reconstructed tomograms at the binning of four were corrected for missing wedge and denoised by IsoNet^65^ version 0.2, applying 35 iterations with a sequential noise cut-off level of 0.05, 0.1, 0.15, 0.2, 0.25 at iteration 10, 15, 20, 25, 30. Thylakoids and pyrenoid tubules were initially segmented using MemBrain-seg^66^, and then imported into ChimeraX^48^ for cleaning and polishing. Rubisco particles were mapped back into the tomographic space based on their refined positions and orientations. The Rubisco model in the segmentation was adapted from the in-cell structure in this study.

### Measurements of pair-wise distance and angle

The distance between adjacent Rubisco particles was calculated according to the coordinates of their centres after the refinement. Paired particles with a centre-to-centre distance shorter than 105 Å were regarded as duplicates and removed. The distribution and local clustering analysis of Rubisco were performed using the software MagpiEM (https://github.com/fnight128/MagpiEM).

## Data availability

Raw tilt series have been deposited in EMPIAR (EMPIAR-12515). The best-resolved (class 12) subtomogram averaging map and corresponding model have been deposited in the EMDB and PDB, with accession codes EMD-52438 and 9HVM, respectively. Subtomogram averaging maps from D4-symmetric classes 7, 10 and 13, and asymmetric class 12 map, have been deposited as additional maps under the same EMDB entry.

## Acknowledgments

We thank Daniel K. Clare for support and David Farmer for help with scripting. We thank Diamond Light Source for access and support of the cryo-EM facilities at the UK National Electron Bio-Imaging Centre (eBIC) (proposal NT29812 and NR21005). The work was supported in part by a grant from the Estate of Louise Yasgour, by the National Institutes of Health (P50AI150481), the UK Wellcome Trust Investigator Award (206422/Z/17/Z), and ERC AdG grant (101021133) and National Institutes of Health grants U54 AI170791 and R01AI178846. This work used Stampede3 at TACC through allocation MCB-170096 from the Advanced Cyberinfrastructure Coordination Ecosystem: Services & Support (ACCESS) program, which is supported by National Science Foundation awards #2138259, #2138286, #2138307, #2137603, and #2138296.

