## supplementary information for "In-cell Structure and Variability of Pyrenoid Rubisco"

### List of Figures

**Supplementary Fig. 1. STA processing pipeline and the main maps produced. a** Classification following two rounds of cleaning "bad" particles. All twenty classes are shown. **b,c** Refinement with C1 (b) and D4 (c) symmetries applied. Refinements that converged and feature overall Rubisco structure are shown. Map numbers correspond to the 20 classes in panel (a) above. The number of subtomograms in each class and maps' resolution are summarized in Supplementary Table 3. **d** The best-resolved map, refined from class 12. ............................. 12

**Supplementary Fig. 2. Fourier Shell Correlation plots for the best-resolved D4 map, refined from class 12.** ............................. 13

**Supplementary Fig. 3. Class 12 map density at key active site fragments.** The relevant fragments are colored red and indicated in each panel, along with the map contour level. Asp473 within the C-terminus is colored blue (e, f). ............................. 14

**Supplementary Fig. 4. Whole domain movements in classes.** The class 12 MDFF model was fitted into the maps of classes 7, 10 and 13. These maps are of lower resolutions, therefore whole domains were fitted as rigid bodies. Domains include the large subunits N-termini (aa 7-148), large subunits C-termini (aa 149-477) and the small subunits. Shown are two large subunits (blue and green) and two small subunits (orange) from each fitted model. The original class 12 MDFF model (grey) is overlayed for reference. ............................. 15

**Supplementary Fig. 9. Creating a hybrid model fitted to the density at both termini. a, c** Visualization of (a) N-terminus and (c) C-terminus of chain E, fitted to the density only at N-terminus. **b, d** Visualization of (b) N-terminus and (d) C-terminus of chain O, fitted to the density only at C-terminus. **e** Superposition of both chains, visualized by secondary structure from residue 7 to 145. **f** Ball and stick visualization of the superposition around residue 103. Superposition of the hybrid large subunit (g) before and after minimization, and (h) all monomers after minimization. ............................. 20

### Detailed methods for real-space refinement of Rubisco coordinates within the cryo-EM map

***Rigid-body docking of Rubisco models into cryo-EM density***

Initially, rigid-body docking was employed to fit the large subunits (PDB: 1GK8[1]) and small subunits (PDB: 1EJ7[2]) of Rubisco structure into the cryo-EM density. Mutations introduced during crystallization were reverted to the wild-type sequence (UniProt ID P00877 for the large subunit and P00873 for the small subunit) using ChimeraX[3].

***Homology modeling of flexible regions of Rubisco***

Flexible regions, namely residue 330 to 339 and 462 to 475 of the large subunits, were generated *de novo* using an iterative procedure with Rosetta version 2024.09[4–7]. Pseudo-D4 symmetry (Supplementary Fig. 13) transformation matrices were created using make_symmdef_file.pl binary in Rosetta, with the matrix details provided in Table S4. One large subunit and one small subunit were considered as a long chain for the symmetry calculation. First, both loop 6 (T330-R339) and C-terminus (W462-L475) of the large subunit were generated using hybridize mover[8] in Rosetta while the pseudo symmetry was imposed. An elec_dens_fast weight of 35 was provided to Rosetta for calculating the energy score that the model would experience in the presence of the cryo-EM density map. The best structure was selected based on Rosetta energy score and visual inspection; the criteria for visual inspection was how loop 6 (T330-R339) and D473 were fitted to the density. Second, part of the C-terminus (W462-T472) of the large subunit that were out of the cryo-EM density were regenerated once again after restraining the distance between D473 Carbon-Gamma and R134 Carbon-Zeta, and H310 Carbon-Gamma, respectively to guide the model into the cryo-EM density. Flat harmonic potentials[9,10] with a standard deviation of 0.25 and tolerance of 1.0, centered at 4.5 and 5.1 Å, for R134 and H310, respectively, were used to model previously observed interactions between these residues[2]. Next, an iterative local rebuilding procedure was applied using CartesianSampler mover in Rosetta. Backbone segments that appeared incorrect based on agreement with local cryo-EM density and their local strain were detected for rebuilding. A fragment length of 7 was used to control the length of the fragments being resampled. New fragments were accepted for replacement when the RMSD between the endpoints of the fragments was below 1.5 Å.

***Molecular Dynamics Flexible Fitting of Rubisco***

Starting from the Rosetta-derived model, protonation states of the titratable groups and coordinates of the hydrogen atoms were added using the PDB2PQR[11] at pH 5.8. Counterions were positioned near the protein using the CIONIZE[12] plugin in VMD[13] with the number of ions determined based on the charged residues present in the Rubisco structure. Ions located more than 15 Å from the protein were removed. The system was then solvated with TIP3P[14] water molecules incorporating a 20 Å padding along each principle axis, resulting in a water box measuring 61× 60 × 72 Å. This solvation was performed using solvate plugin in VMD. A salt concentration of 0.150 M KCl was stablished by addition of bulk ions using autoionize plugin in VMD. The protein structure file for the Rubisco, ions, and water molecules were generated with psfgen plugin in VMD. Secondary structure and chiral restraints were generated using cispeptide, ssrestraint, and chirality plugins in VMD. The system was then minimized using a conjugate gradient algorithm with linear search implemented in NAMD2.15alpha2. Subsequently, Molecular dynamics flexible fitting (MDFF)[15,16] simulations were conducted using NAMD_3.0alpha13[17,18] with the CHARMM36 force field[19]. A time step of 2 fs was employed with the r-RESPA integrator, updating nonbonded interactions every 2 fs, and electrostatic interactions every 4 fs. Long-range electrostatic interactions were computed using the Particle-mesh Ewald (PME) method with a grid spacing equal to 1.0 Å. A cutoff of 12 Å was employed with a switching distance of 10 Å to smoothen the interactions beyond the cutoff. The SHAKE[20] algorithm was applied to restrain all hydrogen bonds. The system temperature was maintained at 300 K using a Langevin thermostat, having a coupling coefficient of 5 $ps^{-1}$. Random initial velocities were assigned to each atom to generate a Maxwell distribution at temperature 300 K. Pressure control was achieved using Nose-Hoover Langevin barostat, at 1.0135 bar, with decay and period of 100 ps and 200 ps, respectively. The backbone heavy atoms were coupled to the cryo-EM density map through a grid-based biasing-potential[21] with a coupling constant of 0.3 au. A harmonic force were applied on all heavy atoms with a spring constant ramping to 2000 $kcal.mol^{-1}.Å^{-2}$ over 1 ns, to transform and overlap the atomic coordinates. Transformation matrices used during the MDFF are provided in Table S5. Secondary structure and chiral restraints were enforced utilizing extraBonds, as implemented in NAMD.

The resulting model was in good agreement with the density as demonstrated by Supplementary Fig. 12a) and supported by the local cross-correlation analysis (Supplementary Fig. 12b-c).

The system obtained from MDFF was then minimized using a conjugate gradient algorithm implemented in NAMD for 40000 steps, ensuring convergence with energy gradient variance falling below 1.0 $kcal.mol^{-1}.Å^{-2}$. During minimization, a coupling constant of 5.0 applied to couple backbone heavy atoms to the cryo-EM density.

***Creating a structure fitted at both termini***

The MDFF-derived model revealed multiple mappings between atomistic coordinates and the cryo-EM density map at the N- and C-termini of the large subunit. While chains fitted well to the density at the C-terminus (Supplementary Fig. 9b), and others fitted at the N-terminus (Supplementary Fig. 9d), no single chain was fitted at both termini simultaneously. To address the latter, two chains were selected based on visual inspection of their fit to the density at each terminus (Supplementary Figs. 9a-d). Their structures were superimposed by aligning their sequences, then fitting their Carbon-Alpha using the matchmaker command in ChimeraX. Only the residues with secondary structure elements from residue 36 to 139 (beta sheets) were selected for superimposing to obtain better superposition (Supplementary Fig. 9e). The superposition of structures allowed us to combine the coordinates of two chains with minimal deviation of atomic positions (Supplementary Fig. 9f).

As a result, a hybrid large subunit was created by combining the residues from 1 to 103 taken from the chain matching the density at N-terminus, while the rest, residues 104 to 475, were taken from the other chain. The hybrid large subunit made by combining these two chains, was subsequently rigid-body fitted to each large subunit in the MDFF-derived model. Finally, the complete Rubisco structure, built from the hybrid model, was subjected to a further relaxation to remove possible steric clashes using NAMD3.0.1.

***Iterative model refinement***

The hybrid model underwent further iterative refinement using a unsupervised[22] and supervised protein refinement tool, applying FastRelax[23–25] protocol in Rosetta. The unsupervised refinement tool identifies regions with low molProbity score[26,27] and refined only those regions while keeping the rest of the structure fixed using MoveMap in FastRelax. Similarly, python and Tool Command

Language (TCL) scripts were developed to identify the regions with poor Local-Cross Correlation (LCC) score and apply FastRelax protocol specifically on those regions (Supplementary Fig. 10).

***Improving the quality of the beta sheets***

Although the protein backbone atoms were well fitted to the density in the refined model, the beta sheets of the model were not predicted properly (Supplementary Fig. 11a-b). In order to enhance the quality of the beta sheets in our model, the transformation matrix from the backbone of beta sheets in a large subunit of PDB 1GK8[1], namely residues 169-171, 199-201, 237-241, 264-268, 290-294, 325-327, 375-379, and 399-40 to the backbone of previously mentioned residues in the refined model were calculated using VMD. Next, these residues were superimposed on the refined model by applying the transformation matrix on them. Then the atomic coordinates of these residues in the refined model were replaced by the coordinates of superimposed residues. Similar procedures were performed on another set of beta sheets, i.e. residues 36-44, 83,89, 97-103, and 130-139.

***Second round of Molecular Dynamics Flexible Fitting of Rubisco***

The coordinates of the hydrogen atoms were added using the PDB2PQR[11] at pH 5.8. The protons added by PDB2PQR to some of the charged amino acids in the structure, i.e. GLU and ASP, were deleted from the structure. The system was ionized, solvated, and underwent an MDFF simulation following the same procedures described earlier. Alternatively, during the MDFF simulation, a harmonic force with spring constant of 4000 $kcal.mol^{-1}.Å^{-2}$ were applied over 0.5 ns on the heavy atoms of the protein. The MDFF were performed utilizing NAMD3.0.1 with CHARMM36m force field[28]. Lastly the model was minimized with NAMD for 40000 steps. The proteins were not coupled to the cryo-EM density during the final minimization. The final structure indicates good agreement with the cryo-EM density map as illustrated by Fourier Shell Correlation (FSC) plot (Supplementary Fig. 12d) and Local Cross Correlation (LCC) between the structure and the map (Supplementary Fig. 12e-f). The beta sheets in the final model are visualized in (Supplementary Fig. 11c-d).

### Supplementary Tables

| Method | Blind milling | Blind milling |
|---|---|---|
| Microscope | Plasma FIB Arctis | Conventional FIB Aquilos 2 |
| Voltage (keV) | 30 | 30 |
| Ion beam source | Argon | Gallium |
| Sputtering coating prior to milling (seconds) | 12 | No |
| GIS coating time (second) | 50 | 30 |
| Bulk milling current | N/A | N/A |
| Milling current | 0.74-2 nA | 0.1-0.5 nA |
| Polishing current | 60 pA | 30 pA |
| Sputtering coating post polishing (seconds) | No | No |
| Fluorescence microscope | iFLM, 100× (TFS) | METEOR, 50× (Delmic) |
| Number of lamellae collected | 37 | 29 |

**Supplementary Table 1. Cryo-FIB lamella preparation.**

| Sample | Arctis lamellae | Aquilos 2 lamellae |
|---|---|---|
| Microscope | FEI Titan Krios G3 | FEI Titan Krios G3 |
| Voltage (keV) | 300 | 300 |
| Detector | Falcon 4i | Falcon 4i |
| Energy-filter | Selectris X | Selectris X |
| Slit width (eV) | 10 | 10 |
| Super-resolution mode | No | No |
| Physical pixel size (Å/pixel) | 1.978 | 1.97 |
| Defocus range (µm) | -2.5 to -4.5, increment 0.3 | -2.5 to -4.5, increment 0.3 |
| Acquisition scheme | Dose-Symmetric, -54º/54º, 2º step, group 3 | Dose-Symmetric, -54º/54º, 2º step, group 3 |
| Total dose (electrons/Å$^2$) | 120 | 120 |
| Number of frames | 6 | 6 |
| Number of lamellae | 37 | 29 |
| Number of selected tomograms/micrographs | 13 | 13 |

**Supplementary Table 2. Cryo-ET data collection.**

| Class | Number of Sub-tomograms | C1 resolution at 0.143 FSC cut-off (Å) | D4 resolution at 0.143 FSC cut-off (Å) |
|---|---|---|---|
| 1 | 12,318 | 24.6 | - |
| 2 | 6,773 | 23.2 | 17.1 |
| 3 | 8,317 | 23.2 | - |
| 4 | 12,275 | - | - |
| 5 | 7,165 | 21.9 | - |
| 6 | 7,797 | - | - |
| 7 | 8,339 | 18.8 | 12.3 |
| 8 | 15,311 | - | - |
| 9 | 9,666 | 23.2 | 17.9 |
| 10 | 12,267 | 19.7 | 13.6 |
| 11 | 7,841 | 28.1 | - |
| 12 | 17,713 | 13.1 | 8.1 |
| 13 | 7,611 | 20.7 | 13.6 |
| 14 | 8,407 | 21.9 | - |
| 15 | 14,197 | - | - |
| 16 | 14,462 | 20.7 | 17.1 |
| 17 | 9,587 | 21.9 | 14.6 |
| 18 | 17,053 | - | - |
| 19 | 11,398 | 17.1 | 15.2 |
| 20 | 6,775 | 22.9 | 17.1 |
| **Total Sub-tomograms** | 215,272 | 148,639 | 104,591 |

**Supplementary Table 3. Number of subtomograms and resolution of the classes.** Class numbers according to the initial classification (Supplementary Fig. 1a) and the number of subtomograms per class are listed in the first and second columns, respectively. Resolution of the same classes refined with C1 or D4 symmetries applied (Supplementary Fig. 1b, c), are listed in the third and fourth columns, respectively.

$$\begin{bmatrix} 1.00 & 0.00 & 0.00 \\ 0.00 & 1.00 & 0.00 \\ 282.20 & 241.06 & 253.39 \end{bmatrix} \begin{bmatrix} -1.00 & 0.01 & -0.01 \\ -0.01 & -1.00 & 0.03 \\ 221.83 & 263.72 & 252.94 \end{bmatrix}$$

$$\begin{bmatrix} -1.00 & -0.02 & 0.00 \\ -0.02 & 1.00 & 0.00 \\ 221.91 & 240.77 & 251.14 \end{bmatrix} \begin{bmatrix} 1.00 & 0.01 & 0.01 \\ 0.01 & -1.00 & 0.02 \\ 282.15 & 263.32 & 251.50 \end{bmatrix}$$

$$\begin{bmatrix} 0.00 & -1.00 & 0.00 \\ 1.00 & 0.00 & 0.00 \\ 241.13 & 221.88 & 253.10 \end{bmatrix} \begin{bmatrix} 0.01 & 1.00 & -0.03 \\ -1.00 & 0.01 & -0.01 \\ 263.11 & 282.45 & 252.72 \end{bmatrix}$$

$$\begin{bmatrix} 0.01 & -1.00 & 0.00 \\ -1.00 & -0.01 & 0.00 \\ 263.34 & 221.79 & 250.79 \end{bmatrix} \begin{bmatrix} 0.00 & 1.00 & -0.03 \\ 1.00 & 0.00 & 0.01 \\ 240.93 & 282.72 & 250.11 \end{bmatrix}$$

**Supplementary Table 4. Rotation matrix used by Rosetta during rebuilding loop 6 and C-terminus.**

$$\begin{bmatrix}1.00 & 0.00 & 0.00 & 0.00\\ 0.00 & 1.00 & 0.00 & 0.00\\ 0.00 & 0.00 & 1.00 & 0.00\\ 0.00 & 0.00 & 0.00 & 1.00\end{bmatrix} \quad \begin{bmatrix}-1.00 & 0.00 & 0.00 & 503.67\\ 0.00 & -1.00 & 0.00 & 504.83\\ 0.00 & 0.00 & 1.00 & 0.09\\ 0.00 & 0.00 & 0.00 & 1.00\end{bmatrix}$$

$$\begin{bmatrix}-1.00 & 0.01 & -0.01 & 505.01\\ 0.01 & 1.00 & -0.01 & 1.45\\ 0.01 & -0.01 & -1.00 & 505.10\\ 0.00 & 0.00 & 0.00 & 1.00\end{bmatrix} \quad \begin{bmatrix}1.00 & 0.00 & -0.01 & 3.24\\ 0.00 & -1.00 & 0.00 & 506.92\\ -0.01 & 0.00 & -1.00 & 505.45\\ 0.00 & 0.00 & 0.00 & 1.00\end{bmatrix}$$

$$\begin{bmatrix}0.00 & -1.00 & 0.01 & 502.61\\ 1.00 & 0.00 & 0.00 & 0.14\\ 0.00 & 0.01 & 1.00 & -1.37\\ 0.00 & 0.00 & 0.00 & 1.00\end{bmatrix} \quad \begin{bmatrix}-0.01 & 1.00 & 0.00 & 0.88\\ -1.00 & -0.01 & 0.01 & 505.14\\ 0.01 & 0.00 & 1.00 & -0.21\\ 0.00 & 0.00 & 0.00 & 1.00\end{bmatrix}$$

$$\begin{bmatrix}0.00 & -1.00 & 0.00 & 504.16\\ -1.00 & 0.00 & 0.00 & 505.20\\ 0.00 & 0.00 & -1.00 & 503.33\\ 0.00 & 0.00 & 0.00 & 1.00\end{bmatrix} \quad \begin{bmatrix}0.01 & 1.00 & -0.01 & -1.07\\ 1.00 & -0.01 & 0.00 & 3.48\\ 0.00 & -0.01 & -1.00 & 506.68\\ 0.00 & 0.00 & 0.00 & 1.00\end{bmatrix}$$

**Supplementary Table 5. Transformation matrices used for imposing symmetry during MDFF simulation.**

### Supplementary Figures

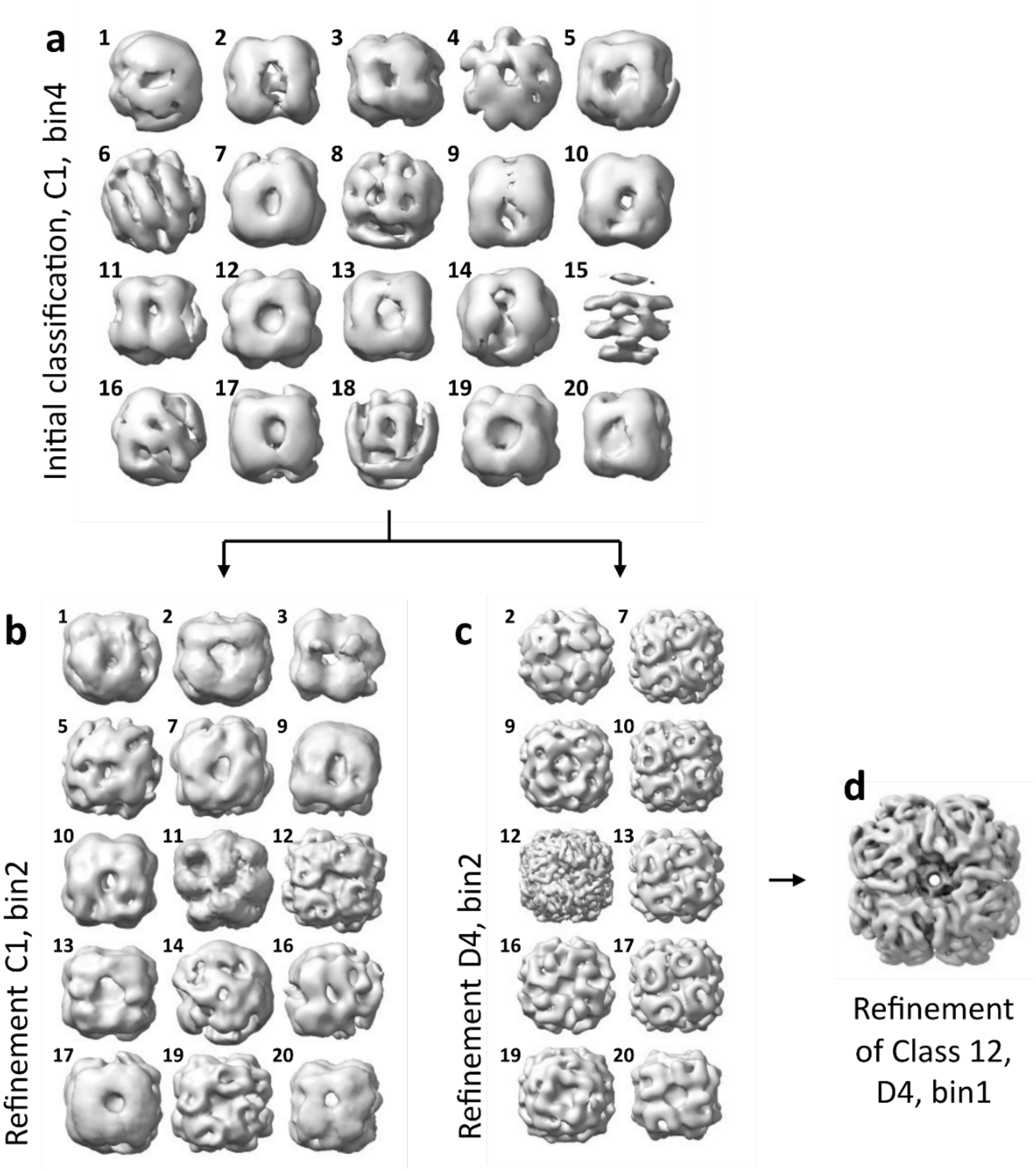

**Supplementary Fig. 1. STA processing pipeline and the main maps produced. a** Classification following two rounds of cleaning "bad" particles. All twenty classes are shown. **b,c** Refinement with C1 (b) and D4 (c) symmetries applied. Refinements that converged and feature overall Rubisco structure are shown. Map numbers correspond to the 20 classes in panel (a) above. The number of subtomograms in each class and maps' resolution are summarized in Supplementary Table 3. **d** The best-resolved map, refined from class 12.

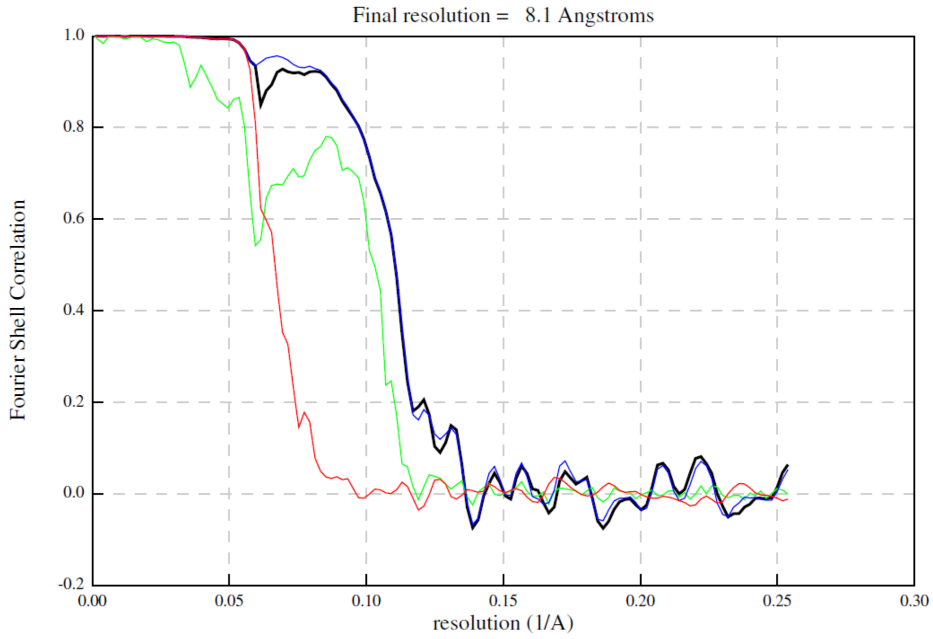

**Supplementary Fig. 2. Fourier Shell Correlation plots for the best-resolved D4 map, refined from class 12.**

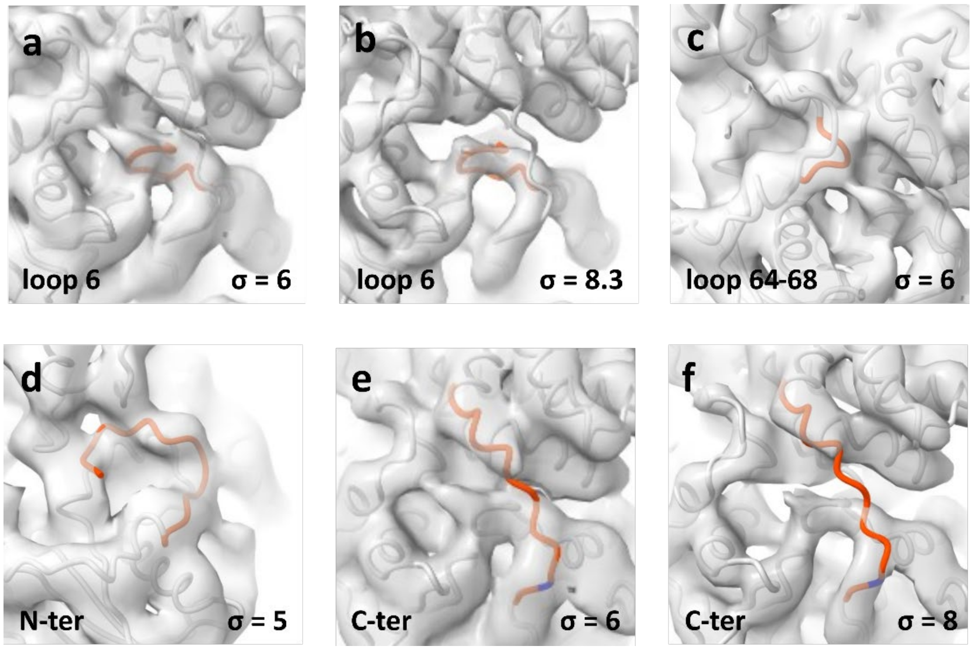

**Supplementary Fig. 3. Class 12 map density at key active site fragments.** The relevant fragments are colored red and indicated in each panel, along with the map contour level. Asp473 within the C-terminus is colored blue (e, f).

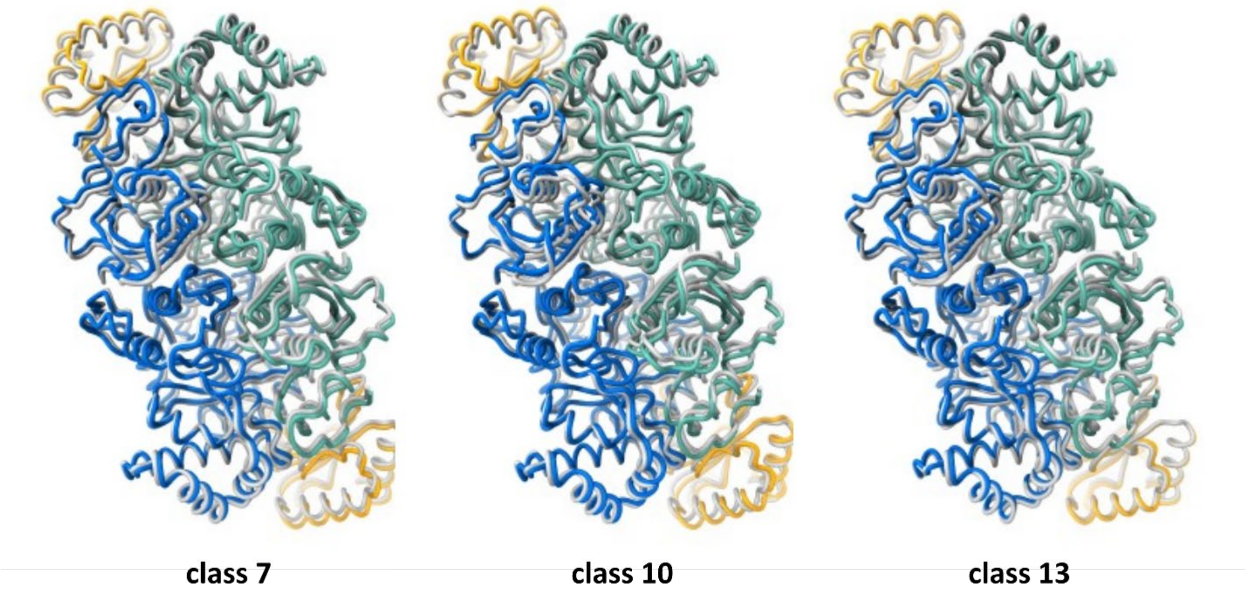

**Supplementary Fig. 4. Whole domain movements in classes.** The class 12 MDFF model was fitted into the maps of classes 7, 10 and 13. These maps are of lower resolutions, therefore whole domains were fitted as rigid bodies. Domains include the large subunits N-termini (aa 7-148), large subunits C-termini (aa 149-477) and the small subunits. Shown are two large subunits (blue and green) and two small subunits (orange) from each fitted model. The original class 12 MDFF model (grey) is overlayed for reference.

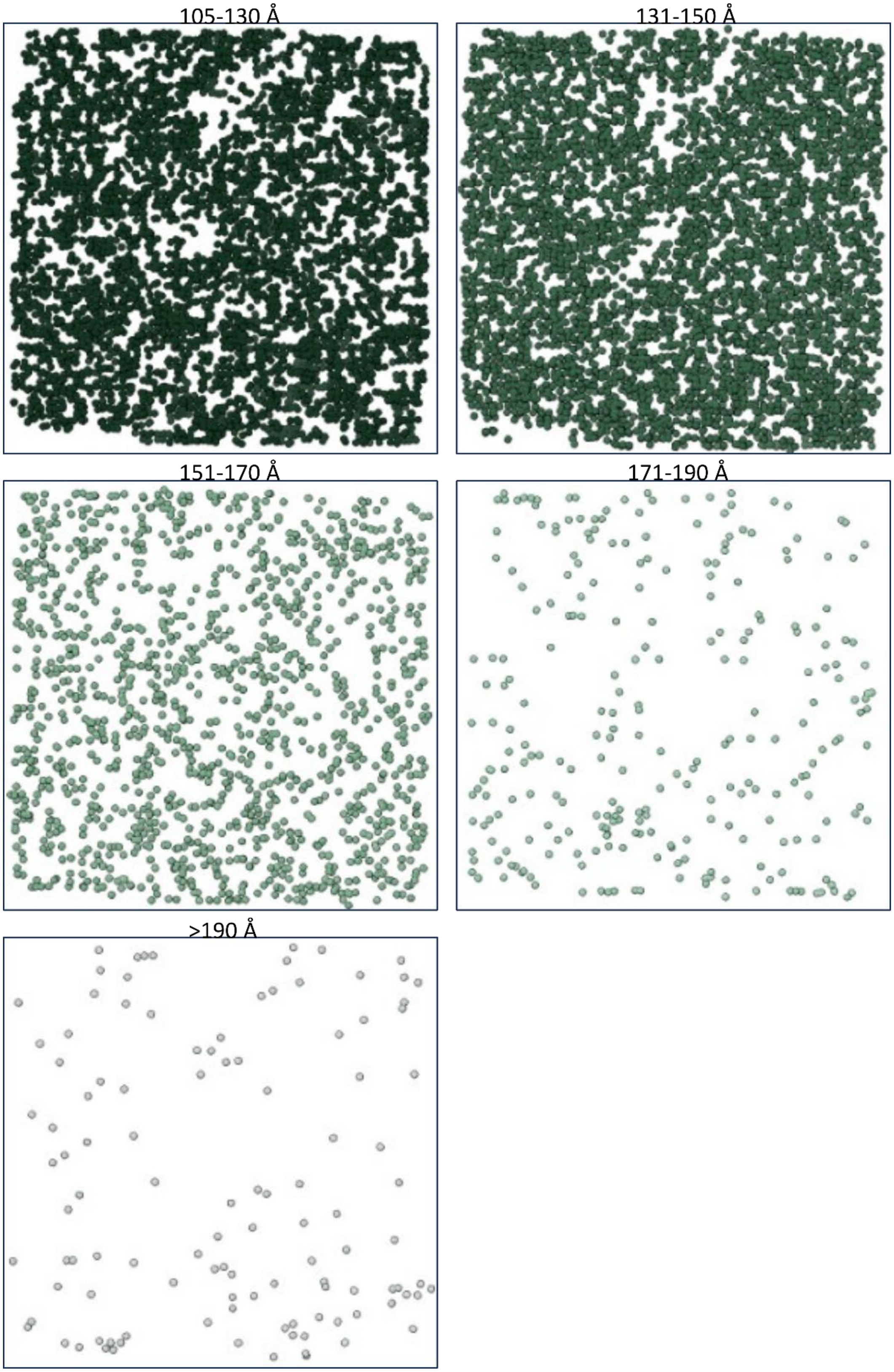

**Supplementary Fig. 5. Individual mapping-back of Rubisco of different pair-wise distances in the pyrenoid from the tomogram in Fig. 5b**

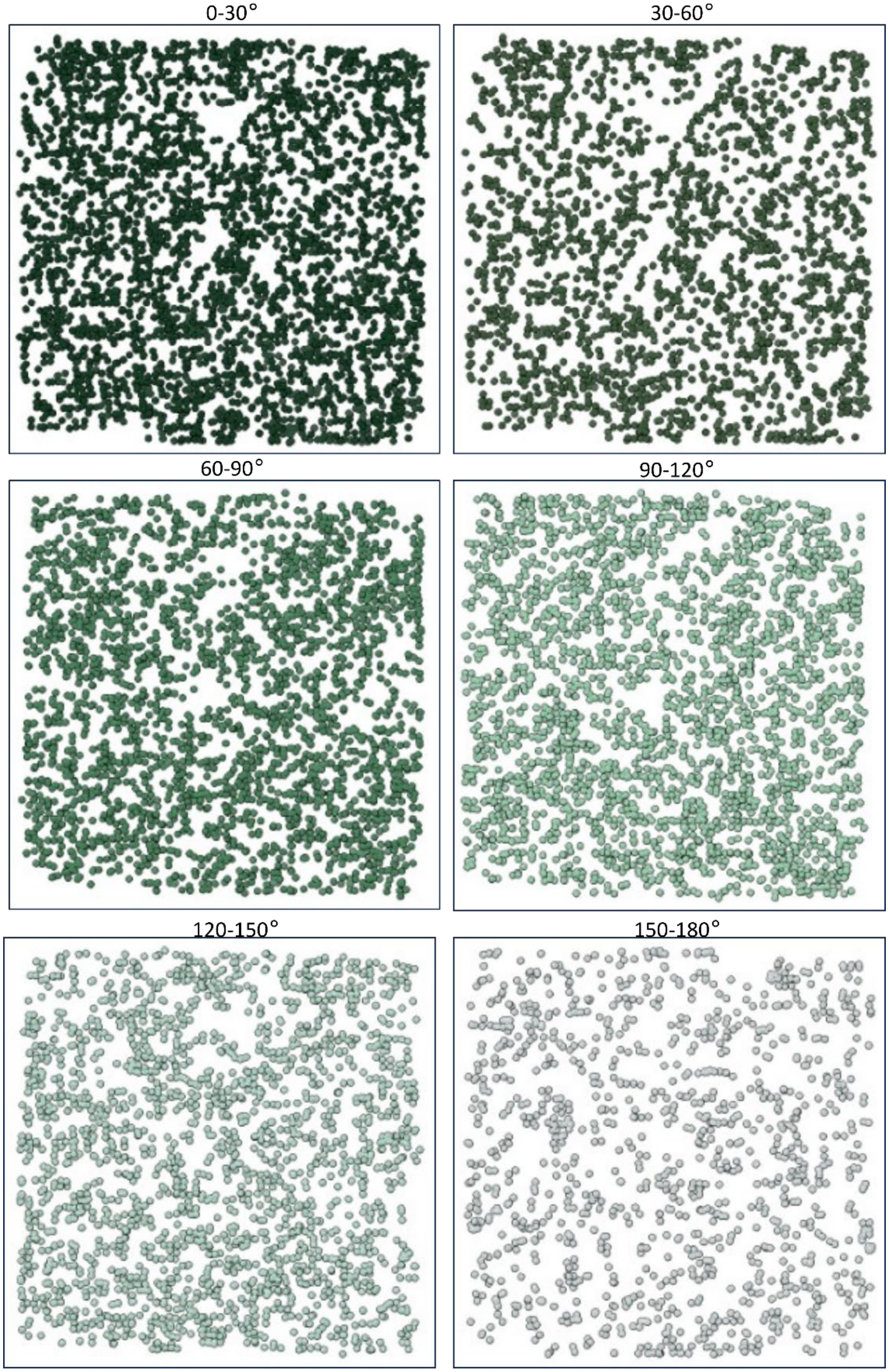

**Supplementary Fig. 6. Individual mapping-back of Rubisco of different paired-wise angles in the pyrenoid from the tomogram in Fig. 5c.**

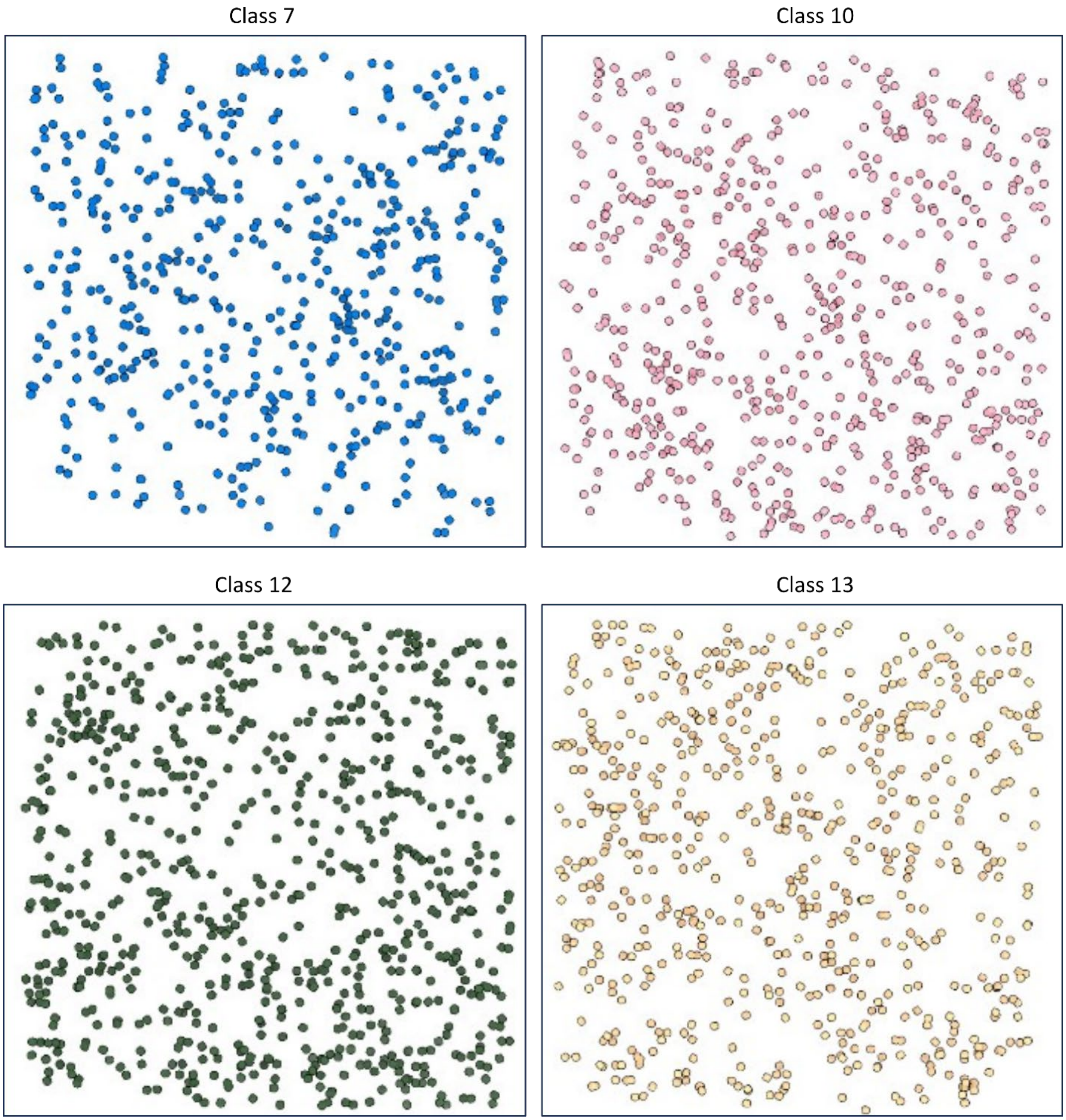

**Supplementary Fig. 7 Individual Mapping-back of Rubisco of different classes in the pyrenoid from the tomogram in Fig. 5d.**

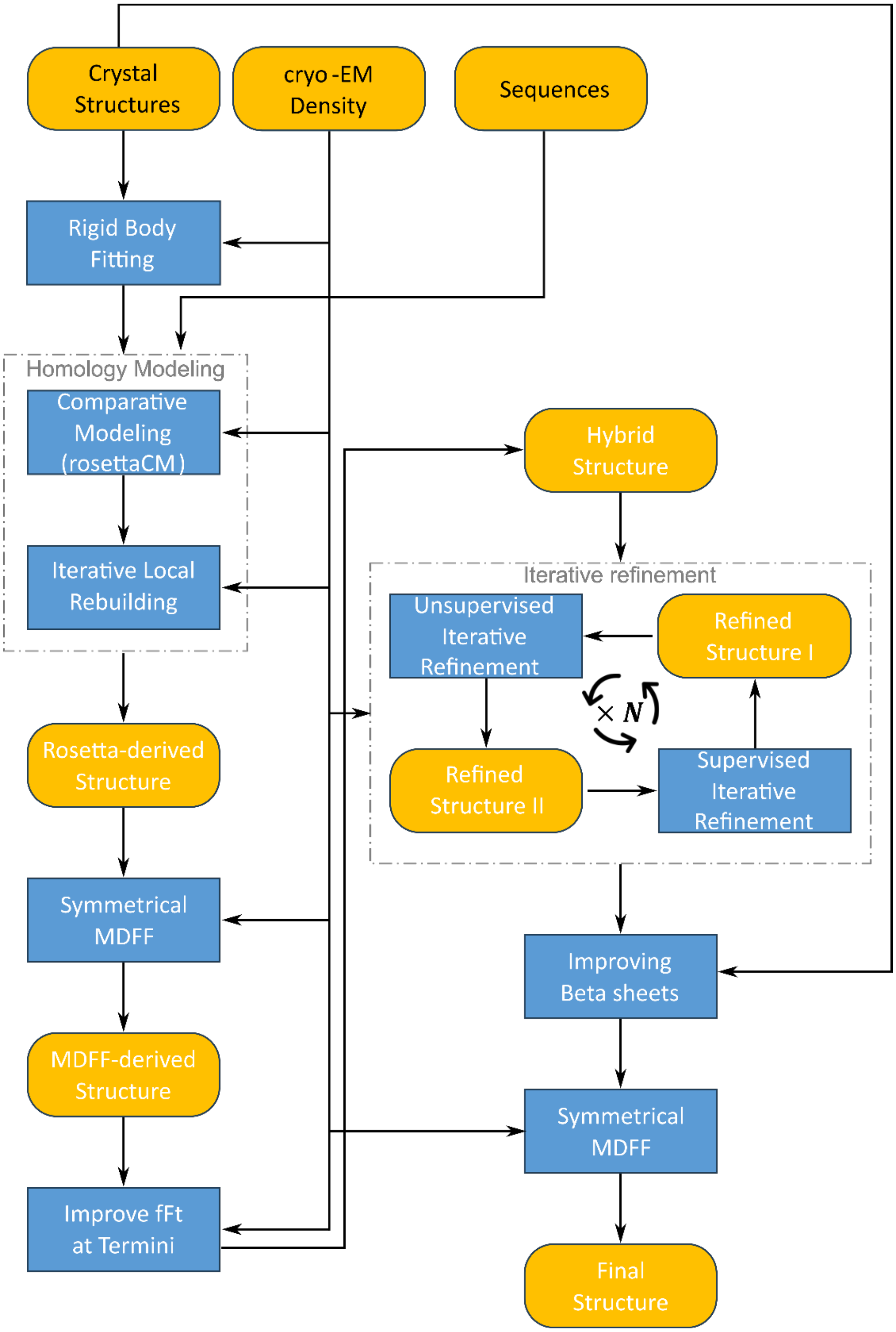

**Supplementary Fig. 8. Flowchart for coordinates refinement, integrative modeling and molecular dynamics flexible fitting simulations.**

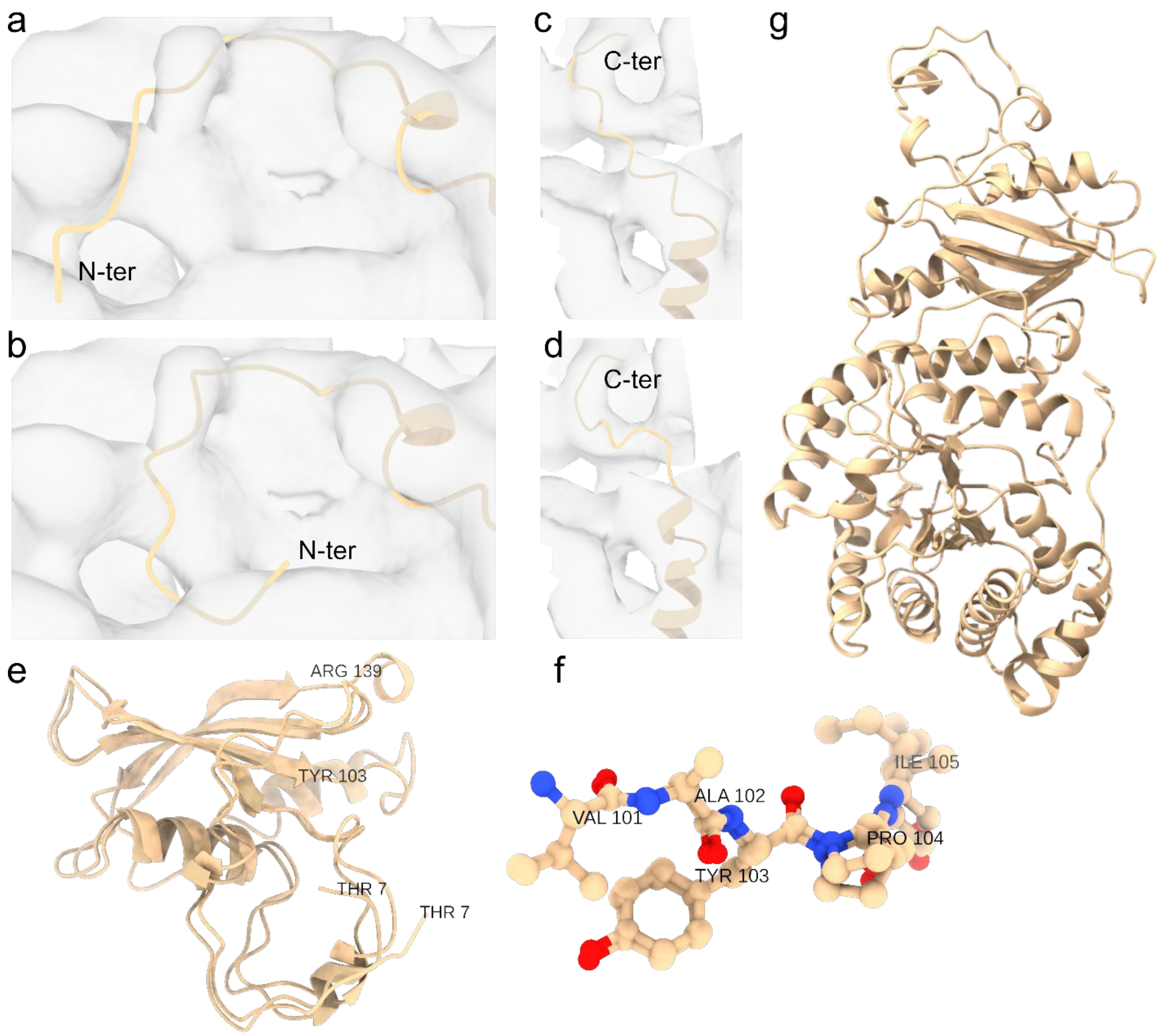

**Supplementary Fig. 9. Creating a hybrid model fitted to the density at both termini. a, c** Visualization of (a) N-terminus and (c) C-terminus of chain E, fitted to the density only at N-terminus. **b, d** Visualization of (b) N-terminus and (d) C-terminus of chain O, fitted to the density only at C-terminus. **e** Superposition of both chains, visualized by secondary structure from residue 7 to 145. **f** Ball and stick visualization of the superposition around residue 103. **g** Superposition of the hybrid large subunit before and after minimization.

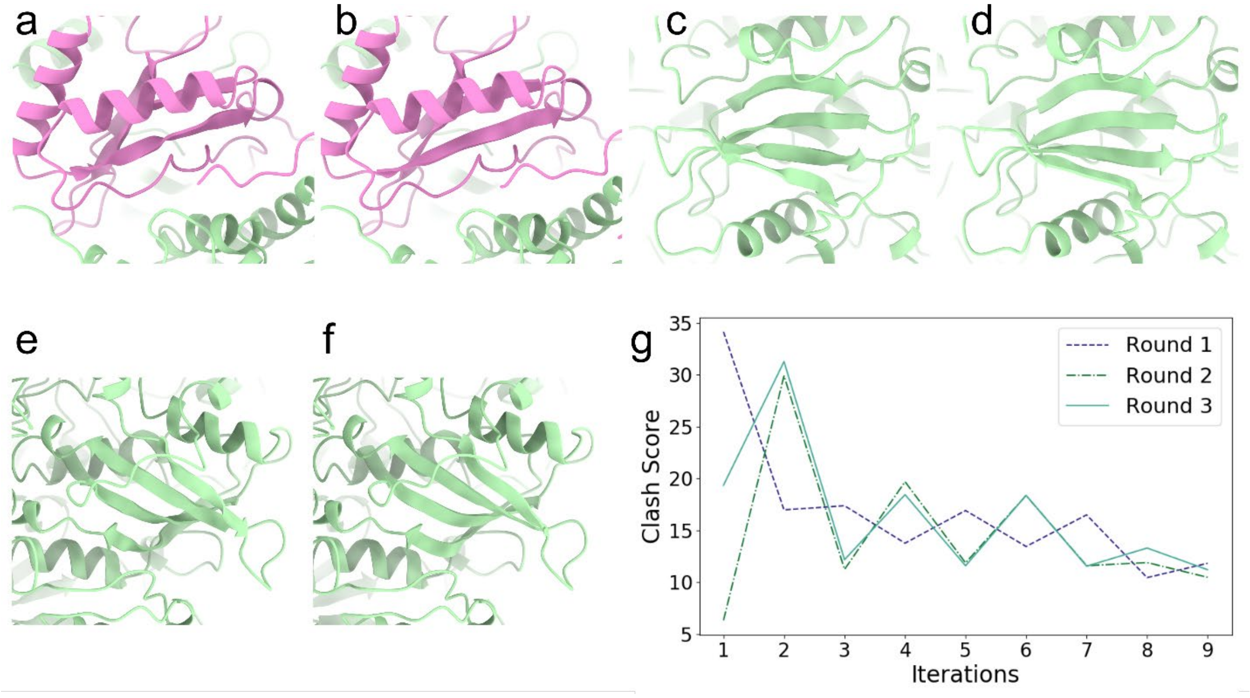

**Supplementary Fig. 10. Model improvement by iterative refinements.** Instances of regions represented by the secondary structure where the refinement tool improved the model, (a, c, e) before and (b, d, f) after applying the refinement tool. **g** Clash score of the structure for iterations of refinement tool.

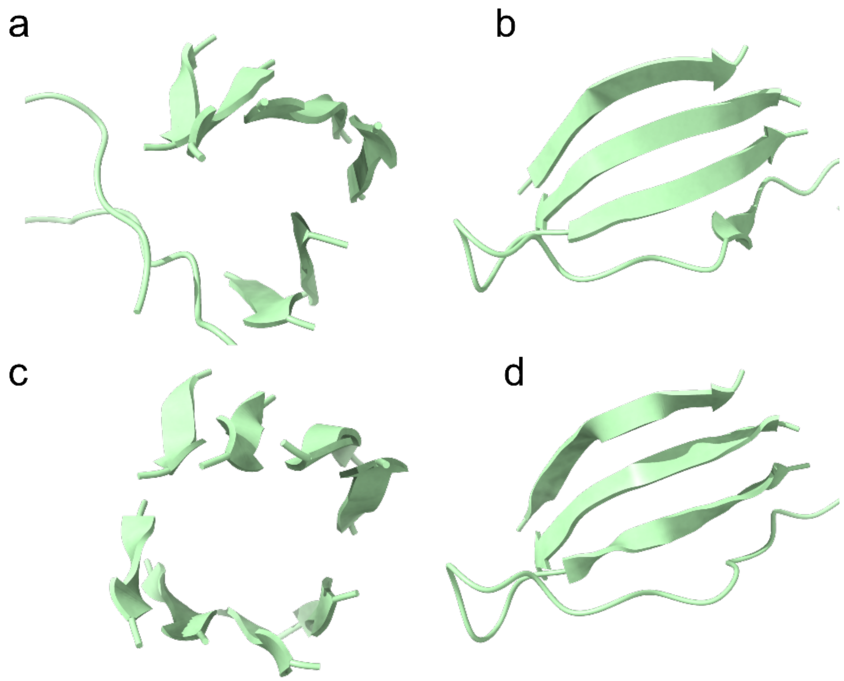

**Supplementary Fig. 11. Improving secondary structures.** Beta sheets in the large subunit in refined model (a, b) and final model (c, d)

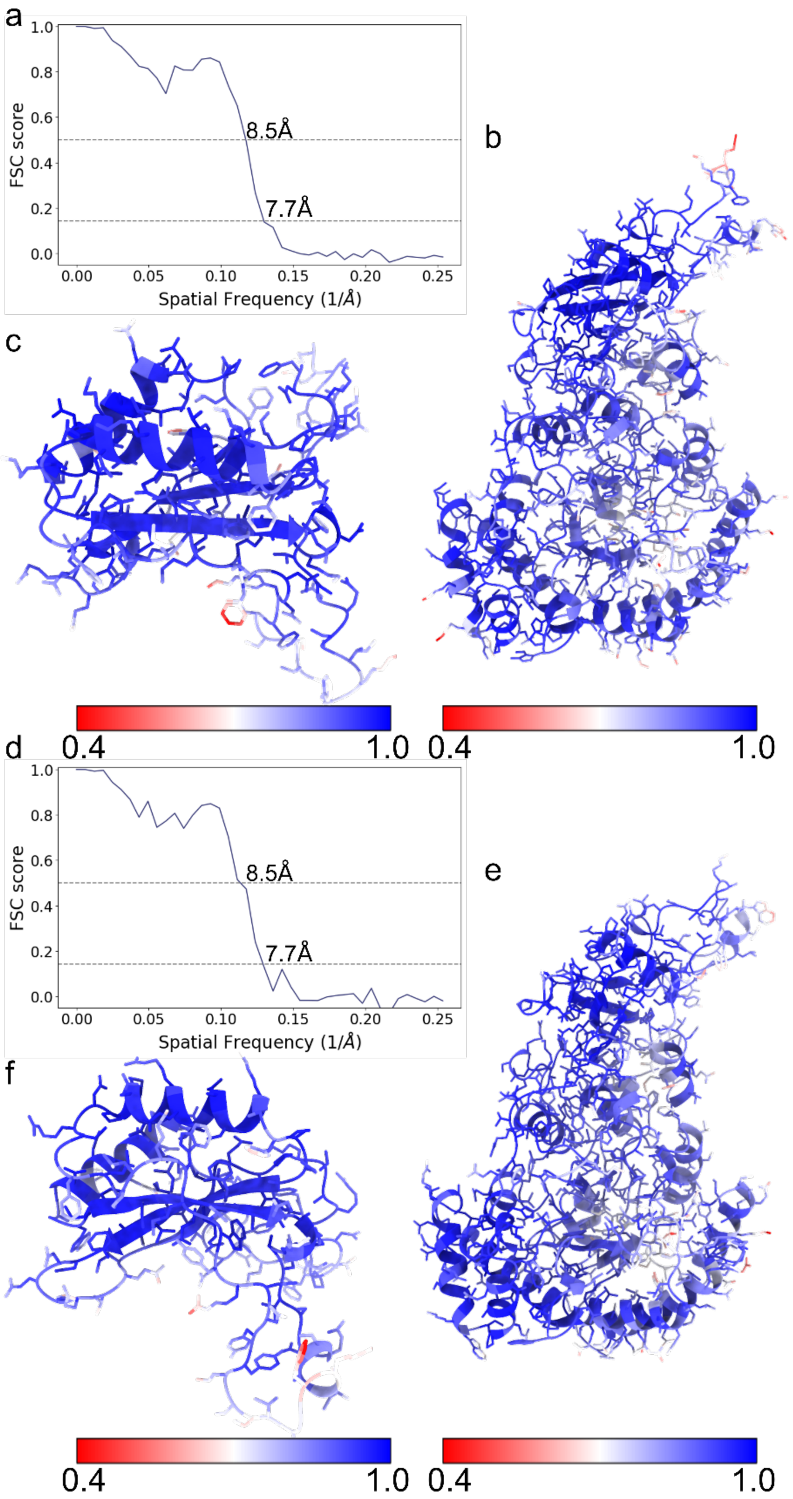

**Supplementary Fig. 12. Quantitative assessment of model agreement to the experimental data. a, d** FSC plot for Rubisco model to density (a) after MDFF simulation and (d) the final structure. **b, c, e, f** Local-Cross correlation (LCC) of the Rubisco structure and cryo-EM density map after MDFF simulation for (b) large and (c) small subunits, and the final structure for (e) large and small subunits.

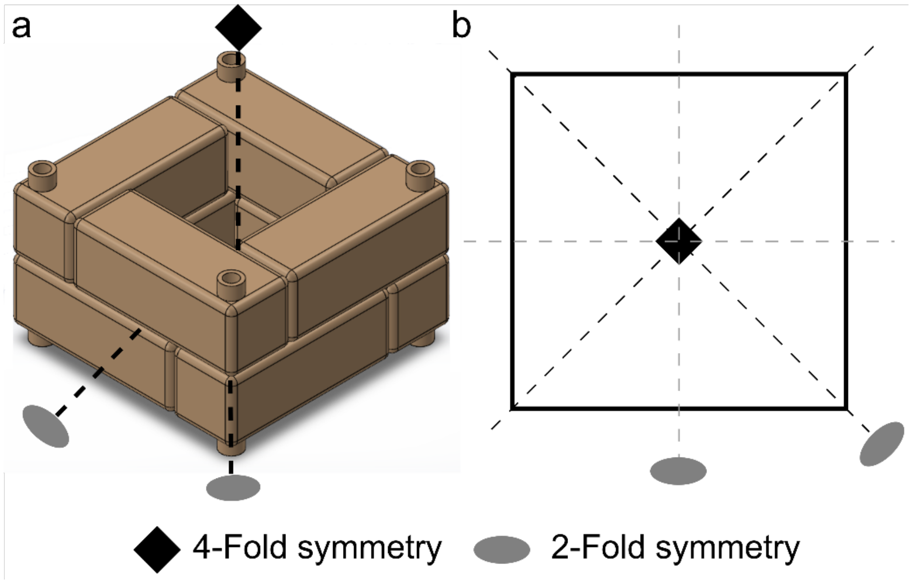

**Supplementary Fig. 13. D4 symmetry:** eight subunits related to each other by one 4-fold axis and two 2-fold axes.
